# A biologically-inspired hierarchical convolutional energy model predicts V4 responses to natural videos

**DOI:** 10.1101/2024.12.16.628781

**Authors:** Michael Oliver, Michele Winter, Tom Dupré la Tour, Michael Eickenberg, Jack L. Gallant

## Abstract

V4 is a key area within the visual processing hierarchy, and it represents features of intermediate complexity. However, no current computational model explains V4 responses under natural conditions. To address this, we developed a new hierarchical convolutional energy (HCE) model reflecting computations thought to occur in areas V1, V2, and V4, but which consists entirely of simple- and complex-like units like those found in V1. In contrast to prior models, the HCE model is trained end-to-end on neurophysiology data, without relying on pre-trained network features. We recorded 313 V4 neurons during full-color nature video stimulation and fit the HCE model to each neuron. The model’s predicted optimal patterns (POPs) revealed complex spatiotemporal pattern selectivity in V4, supporting its role in representing space, time, and color. These findings indicate that area V4 is crucial for image segmentation and grouping operations that are essential for complex vision. Thus, responses of V4 neurons under naturalistic conditions can be explained by a hierarchical three-stage model where each stage consists entirely of units like those found in area V1.

## INTRODUCTION

Visual area V4 is located on the ventral visual processing stream midway between primary visual cortex (area V1) and inferior temporal cortex ^1,2^, and it has been an important target of studies in visual neuroscience ^3,4^. Area V4 appears to represent shape in an intermediate form that is more complicated than the elementary representation found in area V1, but simpler than shape representation in higher-order inferior temporal areas downstream from V4 ^5^. V4 neurons are sensitive to the spatial frequency and orientation of oriented gratings and bars ^6–9^, but polar or hyperbolic gratings elicit much stronger and more selective responses from V4 neurons ^10,11^. Selectivity for polar gratings in V4 suggests that these neurons are tuned for curvature. Indeed, curvature selectivity in V4 has been verified in many studies ^12–18^ and V4 lesions impair curvature discrimination ^19–22^. V4 neurons are also selective for other shape features, such as concave and convex edge elements, sharp protuberances, corners and intersections ^10–14,23^. Some V4 neurons are selective for 3D curvature denoted by stereo ^24–26^, for shape from shading ^27^, or for border ownership ^28^. Many V4 neurons are more selective for texture than for shape ^29–33^. Most V4 neurons are modestly position invariant ^11,34^, though the degree of position invariance depends on the specific features and their distribution within the receptive field ^14,35–37^. Furthermore, many V4 neurons are color-selective ^38–47^. Color selective V4 neurons appear to be concentrated in ’globs’, columnar structures larger than the color-selective ’blobs’ found in V1 ^48^. Most V4 neurons are selective for both shape and color, but some V4 neurons may be selective for only one or the other of these attributes ^49,50^. Because the ventral stream visual pathway is usually associated with object recognition ^2,51^, most studies of V4 have used static, flashed stimuli. However, a few studies that have examined motion selectivity in V4 have reported that roughly a quarter of V4 neurons are directionally-selective ^7,8,52,53^, and that the temporal responses of V4 neurons may differ for shape versus color ^54^. Taken together these various lines of evidence suggest that V4 represents intermediate features of shape and texture, color, and motion. This is consistent with the anatomical position V4 area midway along the ventral processing stream ^1,2^.

Several computational models of area V4 have been proposed previously. The simplest of these describe V4 neuronal tuning within a parameterized space such as spectral features ^56^, curved edges ^12,14^, or color ^47^. These models are primarily descriptive and they do not outline the computational mechanisms underpinning visual processing in V4. Other V4 models have reflected computational theories about how visual representation varies across the primate visual hierarchy ^57–59^. However, these prior models were evaluated using neurophysiology data recorded using simple stimuli or static monochromatic images, so it is unknown whether they accurately predict V4 responses under natural conditions. Because the ultimate goal of any neuronal modeling effort is to predict responses under natural conditions, this is a serious limitation.

To better explain V4 responses to natural images, several recent studies have constructed neuronal models based on features learned by an artificial deep neural network (ADNN) trained to classify static natural images ^60–64^. Because pre-trained ADNNs learn features that reflect the statistical structure of natural scenes, these V4 models tend to outperform those based on parameterized spaces or computational theory. However, this strategy is limited in several ways. First, using a pre-trained model as a source of features for regression may introduce bias due to the specifics of the training stimuli, the architecture of the model and the cost function. Second, fitting a model simultaneously on a population of neurons ^60,61^ may lead to biases in the models caused by correlations between neuronal responses ^65^. In any case, at this time there are no open ADNN models that have been trained on natural movies, so there is no source of temporal features to use for fitting neuronal responses to natural movies. To overcome these problems would require a modeling framework that is based on prior knowledge about the structure of the visual system rather than pre-trained features; which enables one to fit the model to every neuron individually; and which models responses at the fastest time-scale available in the data (i.e., the frame rate of the stimuli for experiments using video displays).

Here we sought to leverage the predictive power of the modern ADNN architecture, while ensuring biological plausibility and avoiding bias in the model’s learned features. Furthermore, we sought to develop an experimental method and a modeling framework that could be used to model the responses of V4 neurons under naturalistic conditions, and at high temporal resolution. To achieve this, we developed an end-to-end-trained, biologically plausible computational architecture called the *hierarchical convolutional energy* (HCE) model. We then used modern artificial neural network training methods to fit this architecture to long-term recordings made from 313 isolated V4 neurons during stimulation with nature full color videos.

The HCE architecture developed here was inspired by earlier attempts to model V4 neurons in terms of known computational mechanisms within the primate visual hierarchy ^57–59^. It consists of a stack of computational subunits designed to encompass operations performed by simple- and complex-cells found in V1, but including no additional mechanisms. In essence, the HCE architecture makes the biologically plausible assumption that areas V2 and V4 both perform precisely the same types of operations on their inputs as those performed in V1. However, the model assumes that, by stacking these linear and nonlinear operations in a hierarchical sequence, area V4 can explicitly represent more complex aspects of shape than the elementary features represented in V1.

Because the computational subunits of the HCE architecture provide an efficient representation of natural images ^66,67^, each model requires orders of magnitude fewer free parameters than those required in popular artificial neural network models, such as AlexNet ^68^. However, the HCE architecture still requires thousands of separate parameters to be estimated for each V4 neuron. Therefore, to ensure that sufficient neurophysiological data were available for this effort we therefore developed a novel long-term recording paradigm for area V4. Long-term recordings were made from Utah recording arrays located chronically in area V4 of two rhesus macaques (see **Figure 1a**). An innovative functional fingerprinting protocol (see **Methods**) enabled us to track single V4 neurons across multiple days, often for several weeks and in many cases for more than a month. The resulting data set consists of recording 313 isolated V4 neurons, probed with up to 1.4 million frames of naturalistic video per neuron.

**Figure 1:**
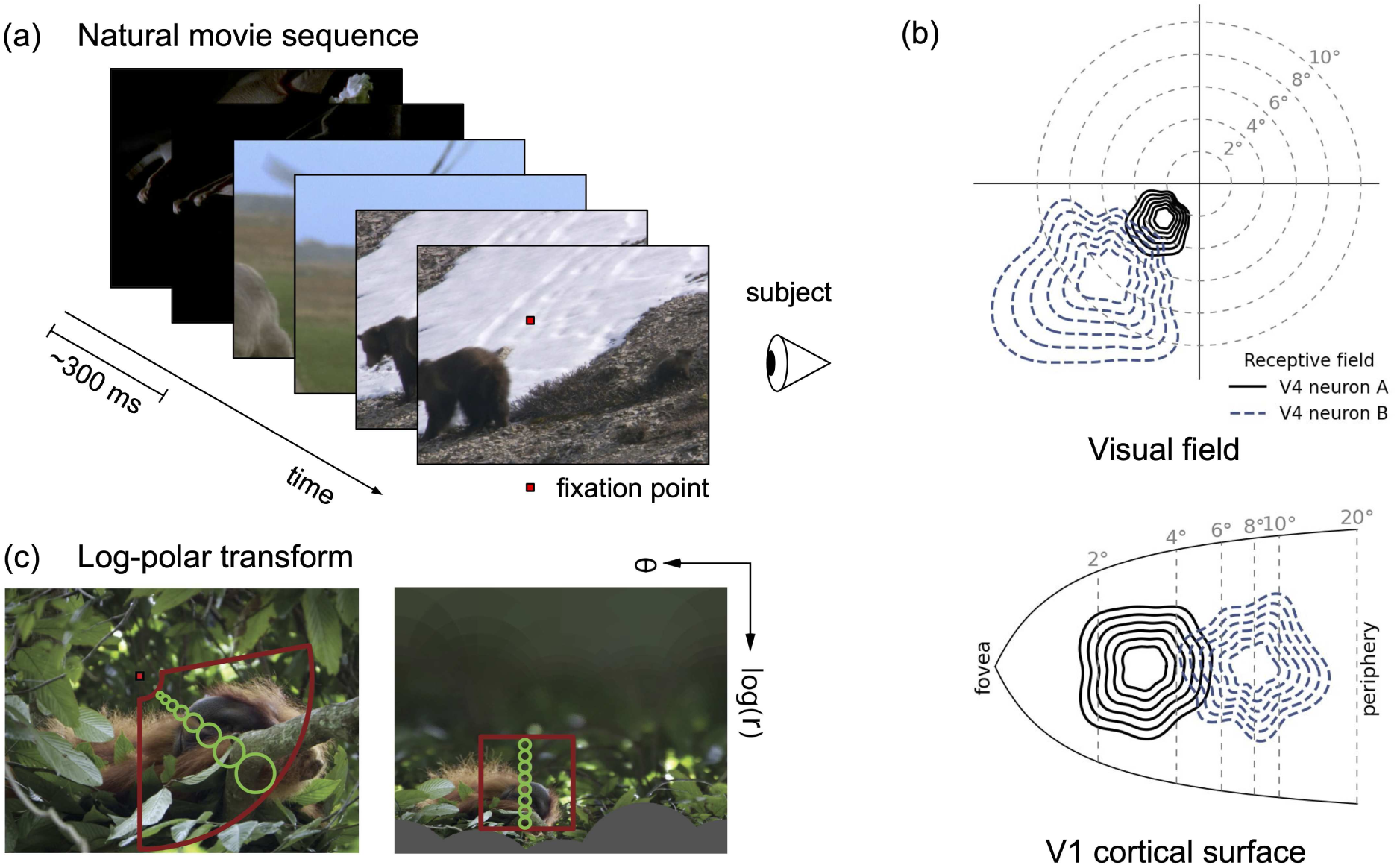
Diagram of the experiment and V4 receptive field organization. **(a)** Diagram of the experiment. To mimic real-world visual conditions, short ∼300ms movie clips were randomly drawn from nature documentaries. Clips were then stitched together to simulate the saccade statistics encountered during natural vision. To more efficiently sample the stimulus space in a limited time frame, the movies were sped up from their native frame rate of 24 Hz to 60 Hz. The videos were presented to animals while the animals were fixating on a small red dot with a black outline. **(b)** Diagram of V4 receptive field organization, adapted from Motter 2009 ^55^. The receptive fields of two V4 neurons (black lined, blue dotted) are projected onto both the visual field (above) and the V1 cortical surface (below). In both spaces, the gray dotted lines denote 2, 4, 6, 8, and 10 degrees of eccentricity. When the receptive fields of the two V4 neurons are projected onto the visual field, the receptive fields are elongated along the radial axis of eccentricity. In contrast, when projected onto the V1 cortical surface, the receptive fields of the two neurons are not elongated along the axis of eccentricity. This suggests that the V4 model should transform the stimulus in order to reduce bias along the axis of eccentricity. **(c)** Log-polar transform. Prior to model fitting, each frame of the movies was transformed from Cartesian space to a log-polar coordinate system. This log-polar transform reflects the cortical magnification that occurs in V1, where pixels near the point of fixation are overrepresented compared to pixels at the periphery of fixation. An example frame in Cartesian space is shown on the left, and the log-polar transformed frame is shown on the right. The pixels contained within the red fan shape in the original image are mapped to the red square in the transformed image. The green circles indicate the receptive field size over different eccentricities. Notice that the varying receptive field sizes are mapped onto equal sizes in the log-polar transformed image.

As noted above, several recent studies of area V4 have sought to model the neural population ^60–64^, rather than modeling single neurons as had been done in prior decades ^12,56–58,69–71^. This likely reflects a shift in emphasis from studies of early vision to studies of higher vision that has occurred over the past several decades: single neuron models are likely most valuable when a visual area represents a relatively simple stimulus properties, so the tuning curves of individual neurons can be well modeled as basis functions in a subspace; population models are likely most valuable when a visual area represents a very high dimensional space, and so the tuning curves of individual neurons are more scattered and random (as happens in compressed sensing ^72^) ^73^. Each approach has its own advantages and disadvantages, and indeed they are complimentary. Because the prior studies of area V4 cited above suggest that it represents features of intermediate complexity, here we chose to model each of the 313 V4 neurons in our sample separately. Fortunately, this was made practical both by the large neurophysiology data set, and by the modest data requirements of the HCE architecture. The resulting models accurately predict the response of their respective V4 neurons to held-out test stimuli, even at the fine 18 ms temporal resolution of the stimulus movies. Finally, to visualize the fit models we synthesized the video sequence that was predicted to elicit the largest possible response from each neuron. These predicted optimal patterns (POPs) confirm prior observations in V4, and they demonstrate that complex pattern selectivity arises in V4 from the combination and recombination of computational mechanisms already known to exist in area V1.

## RESULTS

We performed long-term extracellular recordings (1-108 days) from 313 V4 neurons in two fixating animals during stimulation with naturalistic videos. The final data set consisted of up to 1.4 million frames of video (60 Hz) for each recorded neuron. These data were used to create an explicit, biologically plausible *hierarchical convolutional energy* (HCE) model that accurately predicts responses of single V4 neurons at high temporal resolution and under naturalistic conditions. To interpret the modeling results, the HCE model fit to each neuron was used to predict the spatial, chromatic, and temporal pattern that would be expected to elicit the strongest possible response from that neuron. These predicted optimal patterns were then aggregated across the entire sample in order to quantify the spatial, chromatic, and temporal tuning characteristics of V4 neurons at the population level.

### Prediction accuracy of the HCE Model of V4 Neurons

The HCE architecture is a modular, artificial deep neural network that implements a hierarchy of spatiotemporal filters similar to those thought to exist in visual area V1 (see **Figure 2**; for detailed information on the structure of the model, see **Methods**.) The HCE model takes as input a series of video frames that have been log polar-transformed with respect to the fixation point, and then centered on the receptive field of each V4 neuron under study. The log-polar transform implements the cortical magnification factor inherent in the visual pathway up to and including area V1 (see **Figure 2a**, blue panel). The HCE model itself consists of three separate modules arranged in sequence. The V1 module (see **Figure 2a**, orange panel) implements a collection of spatiotemporal filters like those found in V1 simple and complex cells ^74–77^. This module consists of a complex-valued spatiotemporal convolution operation, followed by complex-valued magnitude and rectifier nonlinearities. The V2 module (see **Figure 2a**, red panel) pools information from V1 and then applies complex-valued spatial convolution and filtering operations similar to those found in area V1 to the incoming signals. The V4 module (see **Figure 2a**, pink panel) pools information from V2. This module is implemented as a spatial receptive field mask that is applied to the incoming signals. All pooling layers are followed by coupled Gaussian dropout layers in order to improve model predictions. The weights within each module and the precise connections between modules are optimized separately for each V4 neuron.

**Figure 2:**
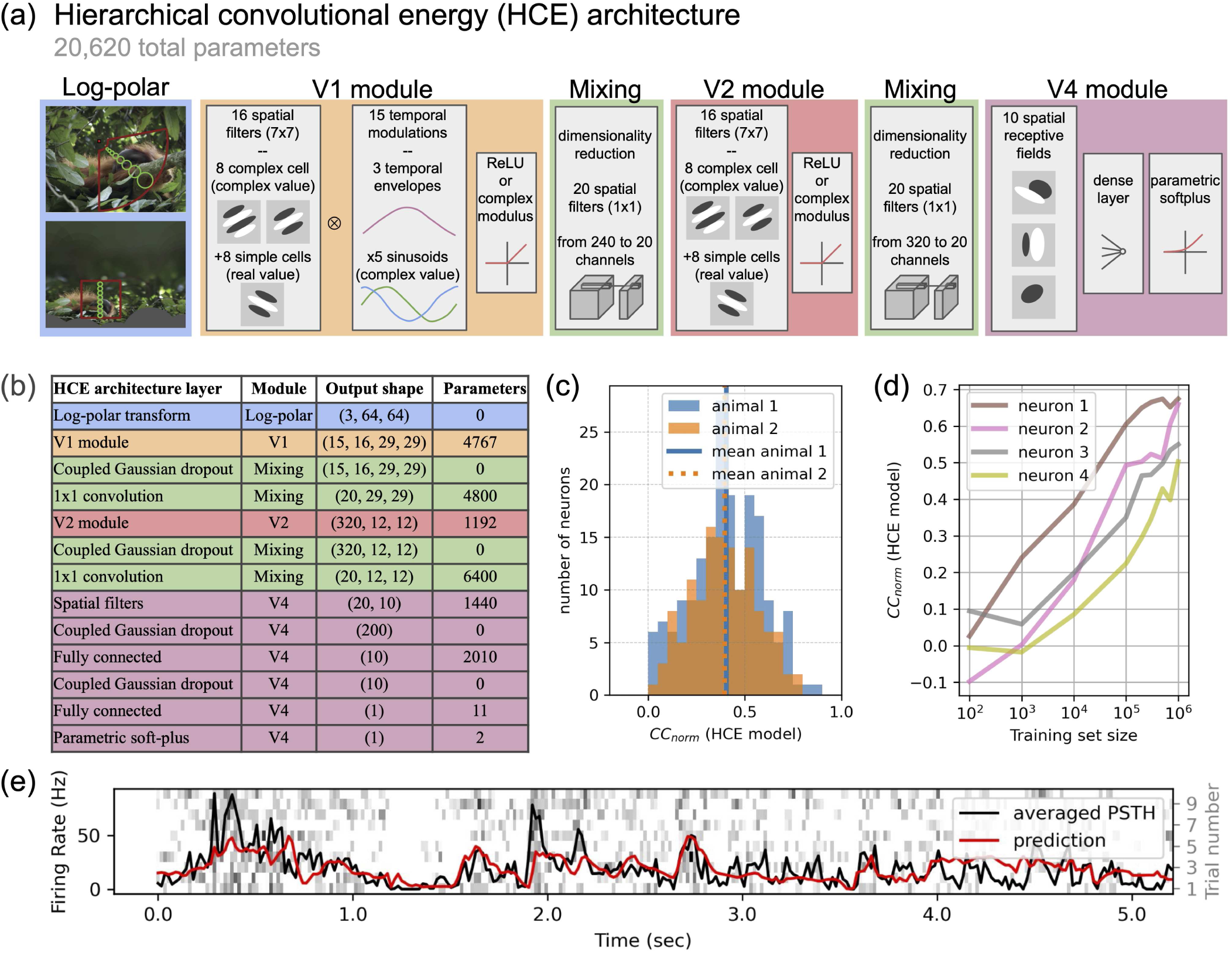
The hierarchical convolutional energy (HCE) architecture and model performance. **(a)** Diagram of the HCE architecture. The HCE architecture is a biologically plausible neural network with three modules that reflect the computational mechanisms that exist in visual areas V1, V2, and V4. A preprocessing stage performs a log-polar transform on the inputs, reflecting the cortical magnification factor (see Figure 1). Signals then pass through a learned bank of spatiotemporal gabor-like filters, reflecting simple and complex cells found in area V1. A mixing module flexibly combines the filter outputs, reflecting neuronal pooling between areas V1 and V2. Signals then pass through a second bank of learned spatiotemporal gabor-like filters, reflecting putative processing in area V2. A second mixing module combines the filter outputs, reflecting neuronal pooling between areas V2 and V4. Finally, signals pass through a third bank of learned spatiotemporal gabor-like filters, reflecting putative processing in area V4. Thus, the HCE model is a hierarchical stack of self-similar V1-like modules, each operating on the output of the earlier layers. **(b)** Output tensor shape and number of parameters in each layer of the HCE architecture. **(c)** Histogram of HCE model performance (CC_norm_) on populations of V4 neurons from the two distinct animals (animal 1 in blue, animal 2 in orange). **(d)** Comparison of HCE model performance on four typical neurons trained with different amounts of training data (shown in logarithmic scale). In all four neurons, larger data sets produce more accurate model predictions. These results suggest that in order to obtain models that predict accurately and that generalize well, researchers should strive to obtain a very large training data set for each recorded neuron. **(e)** HCE model predictions for one well predicted V4 neuron (CC_norm_= 0.74). The figure shows a peri-stimulus time histogram (PSTH) averaged over repeated validation trials (black), and model predictions (red). A visualization of the repeated trials is underlaid in the figure, where each row corresponds to a different trial. The *y* axis on the left marks the 10 different trials. The *y* axis on the right denotes the average firing rate over time.

To determine whether the HCE model provides an accurate description of V4 responses to natural videos, we examined how well the HCE model fit to each V4 neuron predicted neuronal activity in a separate, held out data set. To avoid overfitting, the data recorded from each neuron were first split into a training set that was used to estimate model parameters, and a separate test set that was used to evaluate model predictions. Examination of **Figure 2e** shows that the HCE model provides excellent predictions of the responses of single V4 neurons under naturalistic conditions, even at time scales as fast as the video rate of the monitor. The model performs poorly only in the regions around scene cuts in the naturalistic videos (see **Figure 2e**), where strongly nonlinear response transients in V1 neurons ^78–80^ are propagated up to area V4. Across the sample, the average noise-ceiling-corrected prediction accuracy (*CC_norm_*) of the model at the 16.7 ms frame rate is 0.40 (first quartile 0.29, third quartile 0.53). These predictions are on par with those obtained previously in area V1 studies that used natural images, and in MT studies that used natural movies ^70,77,81^. They also exceed the prediction performance of prior V4 models fit to natural images ^56,81^. Note that correlation measures reported here likely underestimate the true correspondence between V4 neuronal responses and HCE model predictions. V4 responses at the 60 Hz frame rate of the monitor tend to be sub-Poisson, and so violate the Gaussian noise assumption of the correlation coefficient. (Due to the refractory period ^82,83^ cortical neurons are known to be sub-Poisson at short timescales, which strongly biases spike distributions.)

The results shown in **Figure 2** suggest that the HCE architecture provides a good description of the computational mechanisms mediating shape, color and time selectivity in area V4. However, given the complexity of the HCE architecture, it is theoretically possible that the architecture itself might inadvertently overfit the V4 neurons in our sample. To ensure that overfitting did not occur, we developed the HCE architecture using only data from the first animal, freezing the architecture before undertaking analysis of data recorded from the second animal. Then we compared the distribution of model predictions across the two animals. The results show that HCE architecture generalizes across the two animals (see **Figure 2c and 2e**), suggesting that the HCE architecture will also generalize to other animals.

Because we optimized our experiments to provide large data sets, we designed a secondary analysis to evaluate whether our strategy was successful. To do this, we selected a subset of the V4 neurons with very large data sets, and then refit the HCE model to just a fraction of the training data collected in those cells. We find that HCE model performance increases with increasing training set size, and that model performance does not converge even for neurons with over 1 million training data samples (see **Figure 2d**). This suggests that, had our data set been larger, we could have seen a continuous improvement in model performance across our neurons. This highlights the importance of large, naturalistic data sets when training computational models at relatively more central stages of visual processing.

Because several recent studies used a pre-trained artificial neural network (DNN) to model responses of visual neurons ^61–64^, we also compared the performance of the HCE model a model constructed using features drawn from a standard DNN trained on an image classification task (VGG-16 ^84^; see **Supplementary Materials**). to fit a separate model to the data recorded from each of the 313 V4 neurons in our sample. The HCE model performs significantly better than the VGG-F model (see **Supplementary Figure S3**). Thus, the HCE model outperforms a state-of-the-art DNN.

### Using the HCE model to derive the predicted optimal pattern (POP) for each V4 neuron

The architecture of the HCE model contains several nonlinear processing and pooling stages that make it difficult to recover information about spatial, chromatic, or temporal tuning properties of the modeled V4 neurons. This interpretation problem is not unique to the HCE model; both conventional ADNN models ^85–87^ and neuronal models based on ADNN features ^60–64,65^ are notoriously difficult to interpret. In fact, any model that consists of a hierarchical cascade of nonlinear operations presents real challenges for interpretation. To facilitate interpretation of the fit models, we first identified the 14-frame segment of the stimulus videos that had elicited the strongest response from each of the 313 V4 neurons in our sample. This movie clip was then modified iteratively by gradient ascent, until the response of the corresponding HCE model was maximized. The resulting spatio-chromatic-temporal pattern is the stimulus which would be expected, based on the HCE model fit to each V4 neuron, to elicit the strongest possible response from that cell. In this paper we refer to these predicted optimal patterns as POPs (see **Figure 3**).

**Figure 3:**
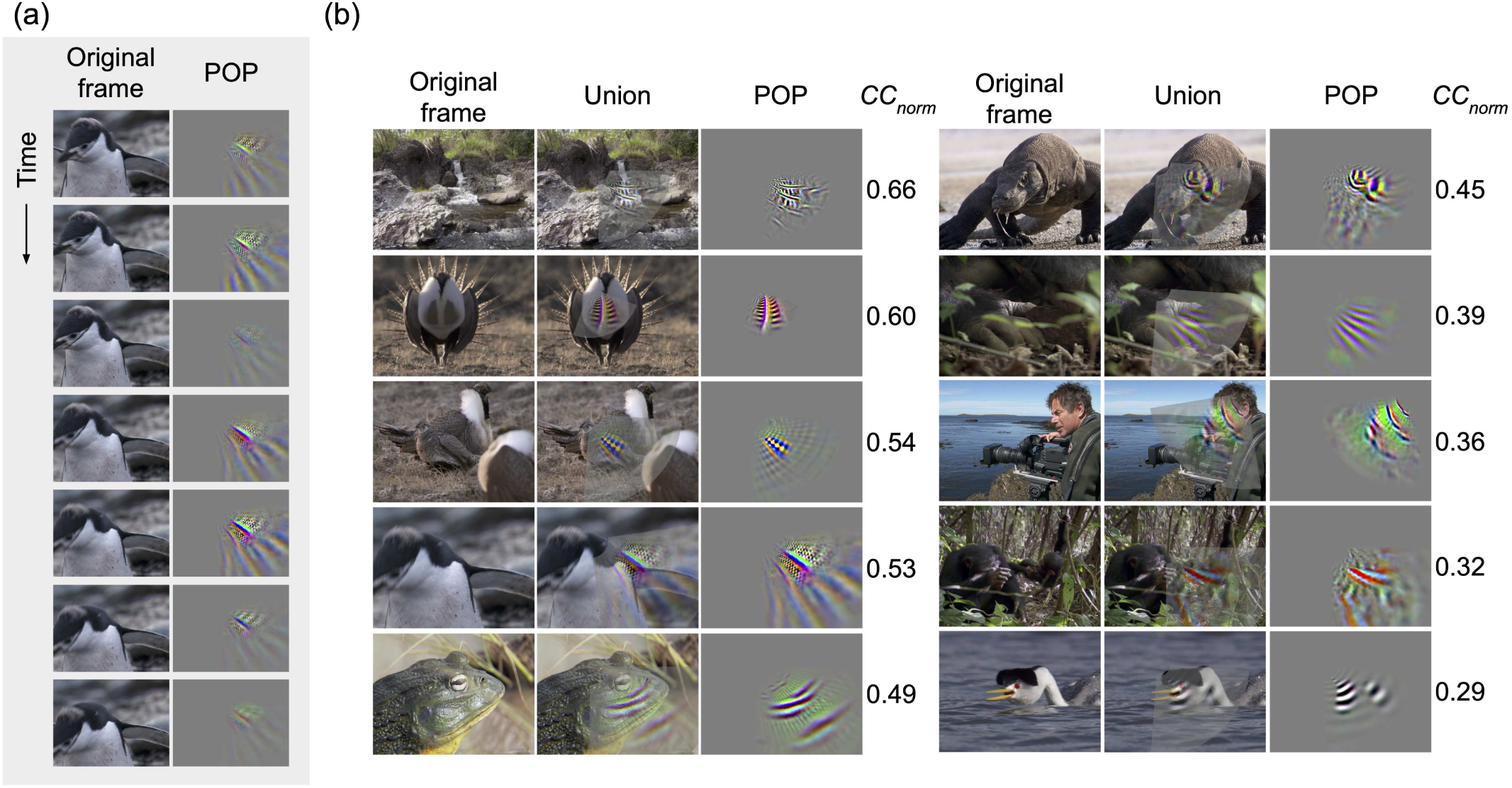
Predicted optimal patterns (POPs) for different V4 neurons. The POPs are synthetically generated stimuli that maximally drive the HCE models trained on the V4 neuron responses. **(a)** Stimulus video that elicited the maximal response from one typical V4 neuron, and the POP for the same neuron. Seven frames from the natural movie stimuli are presented vertically in temporal sequence from top to bottom, and the corresponding POP is shown to the right of each frame. The POP shows which element of the stimulus video elicited the response from the V4 neuron. **(b)** Additional examples of maximally-driving stimuli and corresponding POPs for ten V4 neurons. The left-most frame in each column shows the stimulus video frame that maximally drove the V4 neuron, and the right-most frame in each column shows the POP of the same neuron. The center frame shows a combined visualization, highlighting the correspondence between the POP and the original video frame. For example, the top right triplet shows a neuron with a POP containing tight concentric curves, which correspond to the eye of a Komodo dragon.

### Distribution of spatial tuning properties across the V4 sample

V4 neurons represent shape features that are more complex than those represented in area V1, but less complex than those represented in inferior temporal areas ^6–9,10,11,12,23,88,29,89^. This suggests that V4 neurons are highly specialized shape processing units that combine information about curvature, intersections, and textures to facilitate more complex shape representation in areas downstream of area V4. However, prior studies that examined responses to parametric shape elements may have missed aspects of shape or spatial representation that fell outside the stimulus or model space. To overcome this limitation we sought to develop a data-driven method that could recover all aspects of shape and spatial representation and tuning in area V4. To do this we focused on a spectral analysis of the POPs estimated for each V4 neuron. We first computed a spectral embedding that quantifies the distribution of spatial frequencies in the POPs while marginalizing across color and time. Principal component analysis was used to find the directions of maximum variance in spatial tuning over the V4 sample. A bootstrap procedure was used to identify the most stable spatial PCs.

Applying this procedure to the spectral embeddings recovered from all 313 V4 neurons produced four stable PCs, accounting for 26%, 19%, 12% and 8% of the variance in spatial tuning across the sample, respectively. Projecting the POPs onto the first PC reveals that this PC distinguishes between V4 neurons that are tuned for low spatial frequencies versus those tuned for high spatial frequencies (see **Figure 4a**). The second PC distinguishes between neurons tuned for radial motifs versus those tuned for concentric motifs. The third PC separates neurons tuned for curves differing in curve tangent direction. The fourth PC further distinguishes between neurons tuned to low spatial frequencies versus those tuned to medium spatial frequencies. These results are broadly consistent with prior studies showing that V4 neurons are tuned for spatial frequency ^6–9^ and for non-Cartesian gratings ^10,11^.

**Figure 4:**
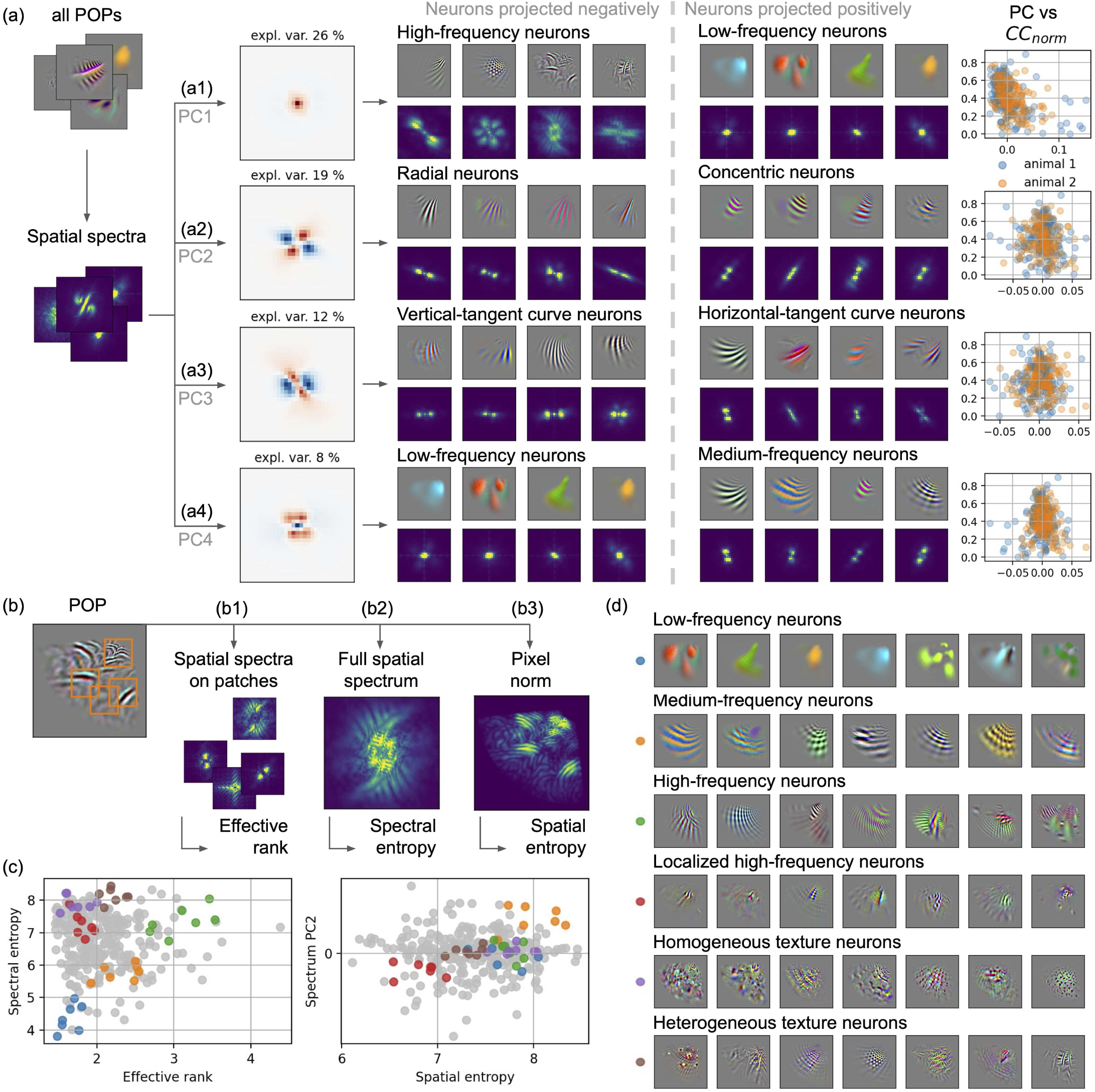
Distribution of spatial tuning over the neuron population. **(a)** To describe spatial tuning, we first consider the spatial spectrum of each predicted optimal pattern (POP). The absolute value of the 2D Fourier transform is used to compute the spatial spectrum, averaging over color channels and over time lags. Then, principal component analysis is used to find the main axes of variation over the neuron population. Bootstrapping was used to determine that the first four principal components (PCs) were stable. For each PC, the four neurons with the maximum negative and positive projections on the PC are shown. Finally, the distribution of neurons along each PC is compared with the neuron prediction accuracy. **(a1)** The first PC separates low-frequency neurons from high-frequency neurons. Note that low-frequency neurons tend to have lower prediction accuracy (see Figure S5). **(a2)** The second PC separates radial neurons from concentric neurons. **(a3)** The third PC separates two sorts of curve neurons, depending on the curve tangent direction. **(a4)** The fourth PC separates low-frequency neurons from medium-frequency neurons. **(b)** To further describe the spatial structures present in the POPs, three additional metrics are computed. **(b1)** First, 1000 patches are extracted from each POP at each time lag, and the spatial spectrum is computed on each patch. Then, a singular value decomposition is used to compute the effective rank, which estimates the number of components necessary to explain the diversity of the spectral patches. A low value indicates high spectral homogeneity over patches of a particular POP. **(b2)** Second, a spectral entropy metric is computed on the full-size spatial spectrum, to estimate the spread of the spatial spectrum over spatial frequencies. A low value indicates high concentration in the spectral plane. **(b3)** Third, a spatial entropy metric is computed on the norm of each pixel, to estimate the spread over pixels. A low value indicates high concentration in the spatial domain. These three metrics and the first four spectral PCs are used to define similarity between neurons. **(c)** Distribution of neurons over four of the seven metrics: the effective rank, the spectral entropy, the spatial entropy, and the second spectral principal component. To show the local homogeneity of this continuous distribution of neurons, six neurons are manually selected from different parts of the distribution. Then, for each selected neuron, the spatial tuning of the selected neuron and its six closest neighbors are visualized. Note that the figures shown are 2D projections from a 7-dimensional embedding **(b4)**. Therefore it is possible that some nearest neighbor clusters may not look like they contain the closest points in the 2D projection, even though they are indeed the nearest neighbors in the higher dimensional embedding. The dotted arrows represent the 2D projection onto spectral entropy and effective rank, and the dashed-dotted arrows represent the 3D projection onto the 2nd spectrum PC and spatial entropy. **(d)** Examples of six local neighborhoods in the proposed neuron distribution. For neuron visualization, the POP is shown at the time lag with the highest norm. The first group (blue) is composed of low-frequency neurons. The second group (orange) is composed of medium-frequency neurons. The third group (green) is composed of high-frequency neurons with strong amplitude modulation. The fourth group (red) is composed of localized high-frequency neurons. The fifth group (purple) is composed of texture neurons that are homogeneous over patches. The sixth group (brown) is composed of texture neurons that are more heterogeneous over patches. Note that these six groups are just samples from a larger distribution. The seven metrics describe a continuously distributed and wide variety of spatial structures that are represented by V4 neurons.

Although the previous analysis provides a broad overview of the most important spatial dimensions represented in area V4, for several reasons the spectral embedding alone cannot provide a complete description of spatial tuning properties within local regions of a neuronal receptive field. Because the spatial spectrum is computed on the entire stimulus, it cannot capture any inhomogeneity in spectral selectivity within the receptive field of a neuron. Furthermore, because the naturalistic movies used in our experiment have a 1/f amplitude distribution, the PCs recovered from the spectral embedding are biased toward low spatial frequencies (see **Figure 4a**). Finally, the spatial spectrum does not take into account the size of the spatial receptive field. Therefore, to obtain a more precise description of spatial tuning properties across our sample of V4 neurons we also examined several additional metrics designed to better reveal the local structure of the spatial receptive field. First, to quantify spectral homogeneity over the POP of each V4 neuron, the local spatial spectrum was computed on 1000 patches extracted from the POP and the effective rank of the distribution of spectra was taken as a measure of inhomogeneity (see **Figure 4b**). A low effective rank indicates that the POP is spatially homogeneous, while a high effective rank indicates that the POP is spatially heterogeneous. Second, the entropy of the full spatial spectrum was used as a measure of the spread of the spectral distribution. Third, the entropy of the pixel norm distribution was used to quantify the spatial spread over pixels, which corresponds to the size of the spatial receptive field.

Combining together the three additional metrics and the PC analysis of the spatial spectral receptive fields described above creates a multidimensional spatial tuning space that describes the distribution of spatial tuning properties of area V4 neurons. Inspection of **Figure 4c** shows that neurons do not fall into discrete clusters within this multidimensional spatial tuning space. Instead, they form a continuous distribution across the space. To provide some intuition about spatial tuning properties in V4, we examined neurons sampled from six different regions distributed across the multidimensional tuning space. For each of these six regions a single neuron was sampled and then this neuron along with its six closest neighbors were plotted. The spectral receptive fields of the resulting 42 V4 neurons are shown in **Figure 4d**. The first set of neurons (first row, blue points) appear to be tuned for low spatial frequencies. The second set of neurons (orange points) appear to be tuned for medium spatial frequencies. The third set of neurons (green points) appear to be tuned for high spatial frequency stripes configured in a complex patchwork of different frequencies, orientations, and colors arrayed across the spatial receptive field. The fourth set of neurons (red points) appear to be tuned for more localized high spatial frequencies. The fifth set of neurons (purple points) appear to be tuned for homogeneous high-frequency textures. The sixth set of neurons (brown points) appears to be tuned for more heterogeneous high-frequency textures. The tuning properties of the six sets of neurons are consistent with the results of the spectral PC analysis reported above, but they demonstrate that the spatial receptive field properties of V4 neurons are far more complicated than can be revealed by any single tuning metric. In fact, most V4 neurons appear to be tuned for spatially inhomogeneous patterns and textures that do not correspond to any of the simple features previously proposed to be represented in area V4.

### Distribution of chromatic response properties across the V4 sample

Area V4 has long been known to contain a high proportion of color-selective neurons ^38,40–42,48,49,90^. To recover chromatic tuning information from the estimated POPs we first constructed a 2D color histogram that quantifies color information in the POPs, while marginalizing across space and time. A principal component analysis was then used to find the directions of maximum variance in chromatic tuning over the V4 sample. A bootstrap procedure was used to identify the most stable spatial PCs.

Applying this procedure to the color histograms recovered from all 313 V4 neurons produced three stable PCs, accounting for 20%, 16% and 11% of the variance in chromatic tuning across the sample, respectively. Projecting the POPs onto the first PC reveals that it distinguishes between V4 neurons that are monochromatic (black and white) and those that are tuned for blue and yellow (see **Figure 5a**). The second PC separates neurons tuned blue and yellow from those tuned to green and magenta. The third PC separates neurons tuned to blue and yellow from those tuned to red and cyan. Thus, these three PCs distinguish between corners of the three dimensional color cube. This confirms many prior reports showing that color-selective V4 neurons are broadly distributed across color space ^47,48,91^.

**Figure 5:**
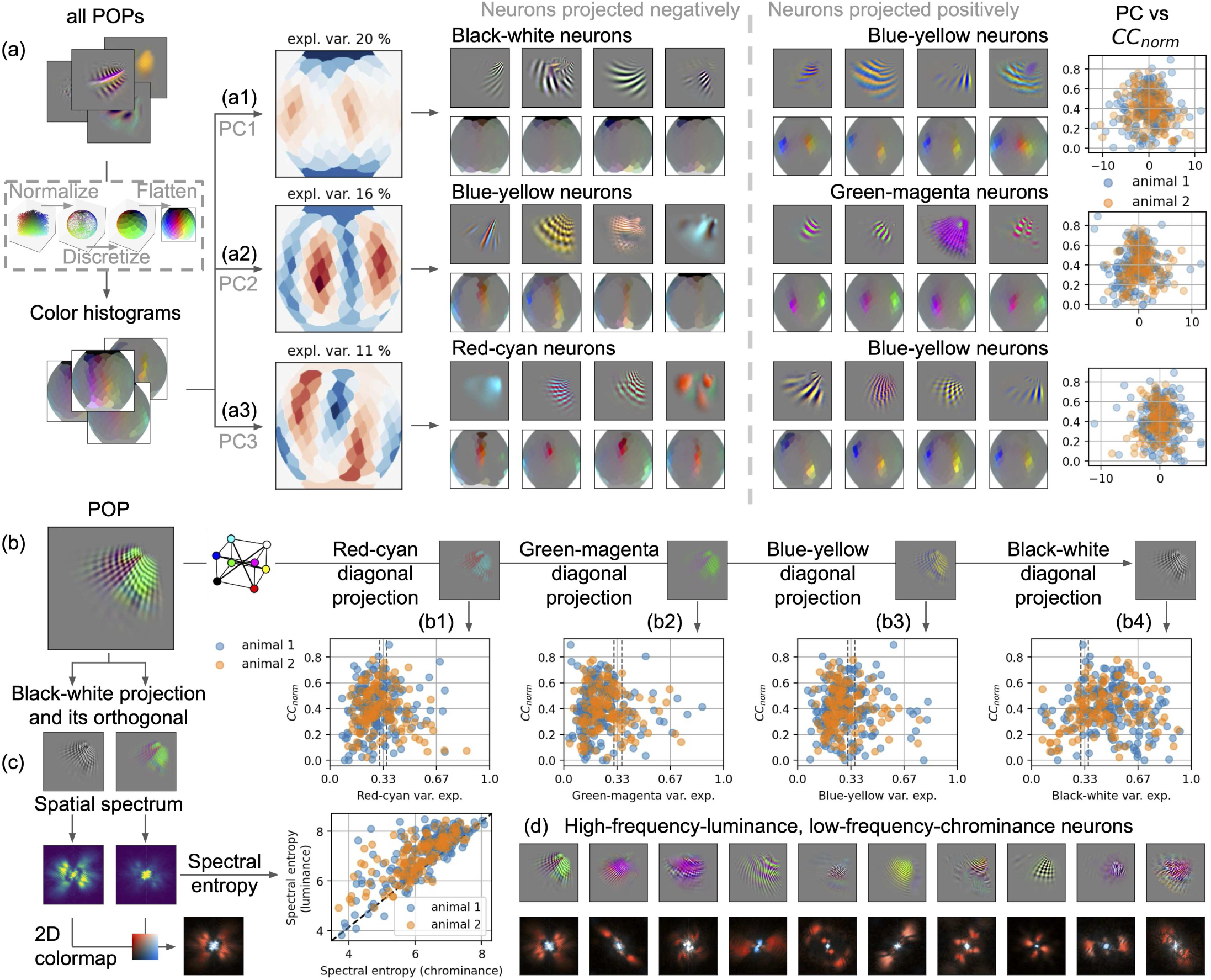
Distribution of chromatic tuning over the neuron population. **(a)** To capture the chromatic tuning on each neuron, an histogram is computed on the color present in each predicted optimal pattern (POP). First, the RGB cube is rotated and normalized to a sphere. Then, the sphere is partitioned into 100 bins of approximate equal size. Finally, the sum of pixels belonging to each bin is computed, weighted by the squared distance of each pixel to the cube center. The obtained spherical color histogram is flatten into a planisphere for visualization. Then, principal component analysis is used on the color histograms to find the main axes of variation over the neuron population. The first three principal components (PCs) were considered stable (bootstrapping). On each PC, the four neurons with the maximum negative or positive projections on the PC are shown. The distribution of neurons along each PC is also compared with the neuron prediction accuracy. **(a1)** The first PC separates black-and-white neurons from blue-yellow neurons. **(a2)** The second PC separates blue-yellow neurons from green-magenta neurons. **(a3)** The third PC separates red-cyan neurons from blue-yellow neurons. **(b)** All three PCs seem to separate one diagonal of the RGB cube from another (black-white, red-cyan, blue-yellow, green-magenta). To systematically compare the four RGB diagonals, the POP is orthogonally projected on each diagonal. Then, the normalized explained variance of each diagonal is computed. A large value indicates that the diagonal explains a large part of the chromatic variance. **(b1-4)** Distribution of the explained variance by each of the four color diagonals, with respect to the prediction accuracy, for both animals. The two vertical dash lines indicate the significance thresholds of chromatic selectivity. A large part of the neuron population (74%) are significantly selective to the black-white diagonal. **(c)** To further describe the prevalence of the black-white diagonal selectivity, the POP is split into a black-white diagonal projection (luminance) and the complementary projection on the diagonal orthogonal plane (chrominance). Then, the spatial spectrum of each projection is constructed, and its spectral entropy is computed. Because the spectral entropy is computed on the normalized spatial spectrum, spectral entropy is independent from the variance explained by each projection. In 298 neurons (95%), the luminance spectral entropy is larger than the chrominance spectral entropy. It reveals that the luminance projection is more spread over high spectral frequencies than the chrominance projection. **(d)** The ten neurons with the largest spectral entropy difference are shown, with at the top the POP at the time lag with the highest norm, and below both spatial spectra superimposed using a 2D colormap. The luminance spectrum is red, the chrominance spectrum is blue, and overlapping frequencies are white. Neurons with a large spectral entropy difference are composed of a high-frequency luminance pattern and a low-frequency chrominance pattern.

The color histogram PCA suggests that although most V4 neurons are tuned for luminance contrast, many of these neurons also have color-opponent response properties. However, this embedding is not optimal for understanding chromatic tuning in V4. The PCs recovered from the color histogram PCs are not necessarily aligned with the cardinal color axes of most theoretical interest. Furthermore, this embedding provides no simple method for quantifying the relationship between luminance contrast and chromatic tuning in V4. Therefore, to better understand potential color-opponent tuning in V4 and its relationship to tuning for luminance we examined two additional metrics of chromatic tuning. First, the POPs were projected orthogonally onto each diagonal of the RGB cube. In this embedding, black/white, blue/yellow and red/green appear at opposite corners of the cube. This embedding provides a simple way to quantify luminance contrast and color opponent tuning by taking the ratio of explained variance of each diagonal with respect to the total variance over the three color dimensions. (Note that because the color space has only three dimensions, the four diagonals are not independent and the sum of the four ratios is greater than one.) Because the HCE model architecture does not favor one color direction over another, neurons that are not color-selective should have color diagonals that each explain about one third of the variance. Neurons that deviate from this pattern are color-selective.

To characterize color selectivity across the sample of 313 V4 neurons, for each neuron, each color diagonal was tested in turn to determine whether it explained significantly more than one third of the chromatic variance (p < 0.03, FDR corrected, permutation test). By this test about three quarters of the neurons are significantly selective for contrast along the luminance axis (233 neurons, 74 %; **Figure 5b**). About one third of the neurons are selective for contrast along the red-cyan axis (91 neurons, 29 %). About one quarter of the neurons are selective for contrast along the blue-yellow axis (81 neurons, 26 %). Finally, about one sixth of the neurons are selective for contrast along the green-magenta axis (49 neurons, 16 %). (Note that these percentages sum to greater than 100% because many V4 neurons are selective for more than one color axis.) These results show that many more V4 neurons are selective to the luminance axis than to any other color axis, a finding that is consistent with previous results ^11,92^.

We also created a new metric to clarify the relationship between luminance and chromatic selectivity in V4. First, each POP was projected separately onto the black-white diagonal (luminance) and onto the orthogonal (chrominance) plane. Then the spectral entropy of both projected embeddings was compared. Spectral entropy measures the concentration of energy over the spectral domain. For example, because low-frequencies are more concentrated than high-frequencies, a high spectral entropy would indicate that a neuron prefers relatively higher-frequency content. According to this metric, the luminance spectral entropy is larger than the chrominance spectral entropy in 298 of the 313 V4 neurons in the sample (95%; see **Figures 5c and 5d**). The difference of spectral entropy is strongly biased toward positive values (range from −0.6 to 3.4; median 0.8). This marked asymmetry in spectral entropy across the sample indicates that V4 neurons are selective for relatively higher spatial frequency luminance modulation, and relatively lower spatial frequency chrominance modulation. This observation is consistent with many prior reports that the primate luminance system is tuned for higher spatial frequencies than is the color system ^93,94^.

### Distribution of temporal response properties across the V4 sample

Because V4 lies on the ventral stream visual pathway ^2,51^ most studies of this area have used static, flashed stimuli to measure V4 tuning for shape or color ^10–13,35,56^. However, a few studies have examined motion selectivity in V4. They have reported that roughly a quarter of V4 neurons are directionally-selective ^7,8,52,53^, and that the temporal responses for the dimensions of shape and color may be different ^54^. Because our study used naturalistic videos that simulated the retinal stimulation that would occur during natural vision, we examined the distribution of temporal response properties across our sample of V4 neurons. We were particularly interested in determining whether V4 neurons are space-time and space-color separable: if the responses of V4 neurons are not space-time separable then the conclusions of prior studies that used flashed images might be in error.

Information about temporal tuning in V4 cannot be recovered directly from the HCE model fit to each V4 neuron, but temporal tuning profiles can be estimated by analysis of the POPs. To recover temporal tuning information we examined the temporal profile of the POPs while marginalizing across space and color. Then, a principal component analysis was used to find the directions of maximum variance in temporal responses over the V4 sample. A bootstrap procedure was used to identify the most stable spatial PCs.

Applying this procedure to the temporal profiles recovered from all 313 V4 neurons produced three stable PCs, accounting for 37%, 22% and 10% of the variance in temporal profiles across the sample, respectively. Projecting the POPs onto the first PC reveals that it distinguishes between V4 neurons that are tuned for relatively more sustained patterns versus those that are tuned for relatively more transient patterns (see **Figure 6a**). The second PC separates neurons tuned to transient patterns depending on their peak’s position in time (i.e. time lags of 2 or 3 frames – 33 or 50 ms – versus time lags of 4 or 5 frames – 67 or 83 ms). The third PC is similar to the first PC, separating neurons tuned to sustained patterns versus neurons tuned to transient patterns. Thus, area V4 neurons appear to fall along a continuum of temporal selectivity, from those sensitive to relatively stationary patterns to those tuned to transient or rapidly changing patterns. This is a common motif seen throughout early stages of the primate visual system. (For analysis of the small differences in temporal responses of V4 neurons found between the two animals, see **Supplementary Materials**.)

**Figure 6:**
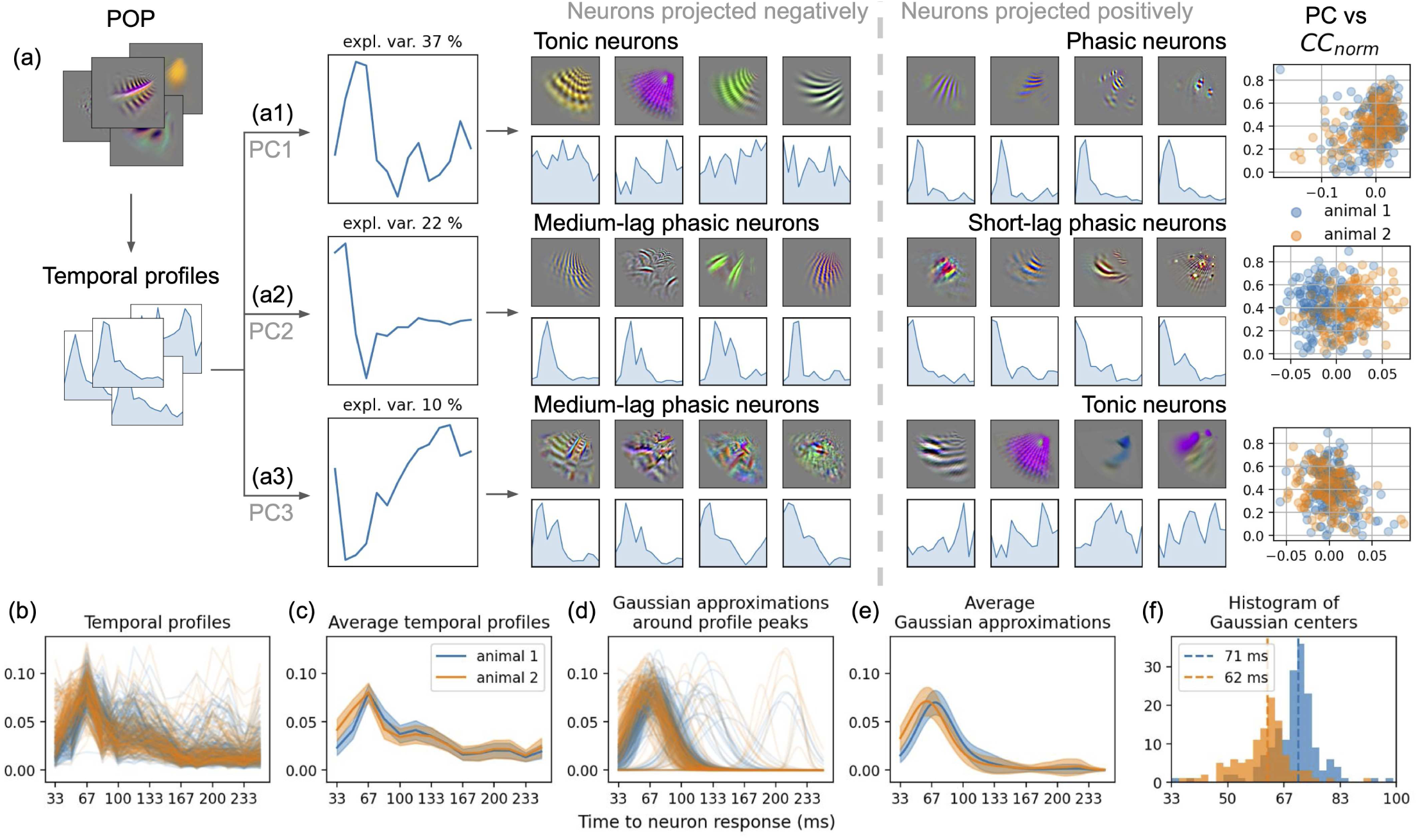
Distribution of temporal tuning over the neuron population. **(a)** First, the norm of the POP at each time lag is computed to create a temporal profile of each neuron. Then, principal component analysis is used to find the main axes of variation over the neuron population. The first three principal components (PCs) were considered stable (bootstrapping). **(a1)** The first PC separates neurons tuned to sustained stimuli over time (tonic neurons) from neurons tuned to brief stimuli (phasic neurons). Note that tonic neurons tend to have lower prediction accuracy (see Figure S5). **(a2)** The second PC separates two sorts of phasic neurons, depending on the peak time lag. Contrary to all other PCs, the distribution along the second PC is different for both animals. **(a3)** The third PC separates medium-lag phasic neurons from tonic neurons. **(b)** To further describe the difference between both animals revealed by the second PC, the temporal profiles were compared between animals. **(c)** Average temporal profile per animal. The shaded areas correspond to one standard deviation around the average. Neurons from the first animal seem to have a peak later in time than neurons from the second animal. However, the 14 time lags give a temporal sampling too coarse to separate the two animals. To circumvent this issue, the temporal profile peaks were approximated with Gaussian functions. **(d)** Gaussian approximations for all neurons, fitted on a window of 5 lags centered on the temporal profile peak. The Gaussian fits give continuous approximations of the peaks, from which a finer peak position can be estimated. **(e)** Average Gaussian approximation per animal. **(f)** Histogram of Gaussian centers below 100 ms. The median peak positions were at 71 ms for the first animal, and 62 ms for the second animal. This 9 ms difference (p < 10^-12^, t-test) could be due to a difference in subject age, in subject neural anatomy, or to a difference in sensor location within the V4 area (assuming different locations have different temporal selectivity).

### Separability of spatial, temporal and chromatic tuning

Many studies of V4 neurons focus specifically on spatial, chromatic or temporal tuning alone, often varying stimuli along the dimension of interest while holding the other dimensions constant. Although this is an efficient way to measure tuning it assumes separability between the three domains, and there is no guarantee that the tuning properties recovered this way will accurately predict tuning under naturalistic conditions where these domains may interact ^95^. We therefore sought to use the rich data acquired here to investigate whether spatial, chromatic, and temporal tuning are pairwise linearly separable across our V4 sample.

To quantify space-time separability we first created a metric that describes separability in pixel space. For each neuron, a singular value decomposition (SVD) was applied over all 14 time lags of the POP ^96^, and the effective rank was computed on the singular values. The effective rank ^97^ is a continuous metric that quantifies the number of components required to reconstruct the spatial pattern of the POP over time. Here, the effective rank takes values in the interval [1, 14]. According to this metric an effective rank close to one indicates that a neuron is space-time separable, while larger values indicate that a neuron is not space-time separable. The distribution of the pixel-based effective rank across all 313 V4 neurons in our sample is shown in **Figure 7b1**. Inspection reveals that only a small minority of neurons have an effective rank close to one (16 neurons out of 313 (5%) are below 1.5), and most effective rank values are between 1.5 and 4.1 (5th and 95th percentiles). Thus, most V4 neurons are not space-time separable.

**Figure 7:**
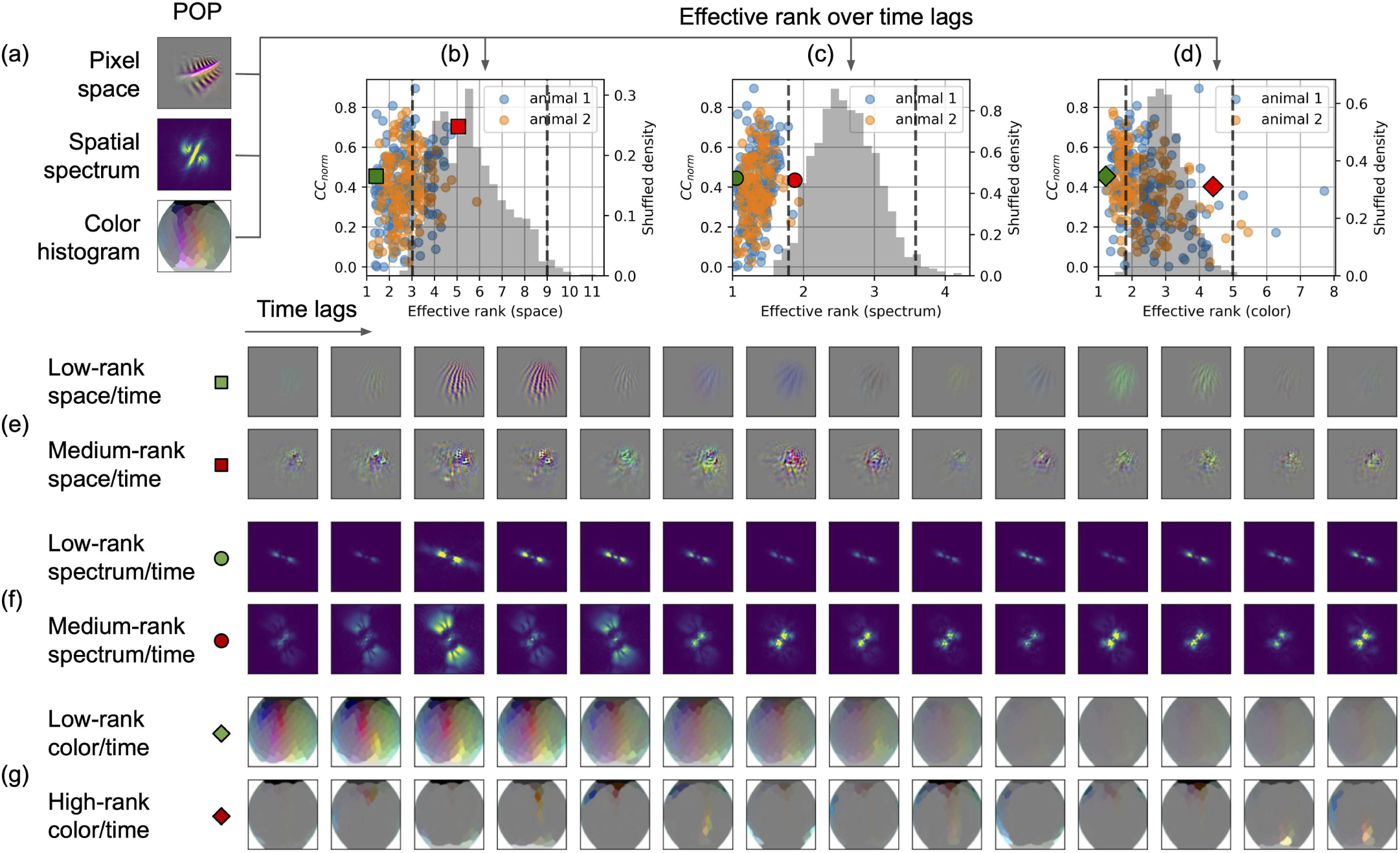
Separability of space, time, and color. To describe the different temporal structures present in each POP, the rank of different space-time and color-time embeddings is computed. **(a)** Three embeddings are considered: the original POP in RGB pixel space, the spatial spectrum, and the color histogram. Each embedding is computed independently on each time lag of the POP. Then, a singular value decomposition is computed over lags to extract the components that are used over multiple lags. Finally, the effective rank is computed based on the explained variance of each component. A low value indicates that a small number of components are repeated over lags. **(b1-3)** Distributions of effective rank (horizontal axis) for all neurons, with respect to the performance of the neuron (*CC_norm_*, left vertical axis). A null distribution (gray, right vertical axis) is computed by shuffling POPs lags across neurons, and computing the same effective rank. The significance thresholds are shown with dashed vertical lines (p < 0.025, FDR corrected). **(b1)** Distribution of effective rank for the original embedding in RGB pixel space. All neurons have a lower effective rank than the null distribution median. However, many neurons do not have an effective rank significantly lower than the null distribution. This result indicates that although POPs have some amount of redundancy over time lags, they do not all have a low-rank structure in pixel space. Examples of a low-rank neuron and a medium-rank neuron are shown in **(c1)**, with the respective positions in the distribution indicated with the green and red squares. **(b2)** Distribution of effective rank for the spatial spectrum embedding. Many neurons have an effective rank significantly lower than the null distribution. This result indicates that many POPs have a low-rank structure in the spatial spectral embedding. Examples of a low-rank neuron and a medium-rank neuron are shown in **(c2)**, with the respective positions in the distribution indicated with the green and red circles. **(b3)** Distribution of effective rank for the color histogram embedding. Some neurons have an effective rank significantly lower than the null distribution, and some other neurons have an effective rank significantly higher than the null distribution. This result indicates that some POPs have a low-rank structure in the color histogram embedding, while other neurons have a high-rank structure. Examples of a low-rank neuron and a high-rank neuron are shown in **(b3)**, with the respective positions in the distribution indicated with the green and red diamonds.

To investigate spectrum-time separability, we created a separate metric based on the spatial spectrum of each POP. For each neuron, the spatial spectrum of the POP was computed at each of the 14 time lags. Then, an SVD was applied over all 14 time lags of the spatial spectrum, and the effective rank was computed on the singular values. An effective rank close to one indicates that a neuron is spectrum-time separable, while larger values indicate that a neuron is not spectrum-time separable. The distribution of the pixel-based effective ranks is shown in **Figure 7b2**. Contrary to the pixel embedding, the spectral embedding leads to effective ranks close to one (260 neurons out of 313 (83%) are below 1.5), and most effective rank values are between 1.1 and 1.6 (5th and 95th percentiles). Thus, most V4 neurons are spectrum-time separable.

Finally, to quantify color-time separability across our sample of V4 neurons, we created a separate metric based on the color histogram of each POP. For each neuron, the color histogram of the POP was computed at each of the 14 time lags. Then, an SVD was applied over all 14 time lags of the color histogram, and the effective rank was computed on the singular values. An effective rank close to one indicates that a neuron is color-time separable, while larger values indicate that a neuron is not color-time separable. The distribution of the pixel-based effective ranks is shown in **Figure 7b3**. Inspection reveals that only a minority of neurons have an effective rank close to one (30 neurons out of 313 (9.6%) are below 1.5), and most effective rank values are between 1.4 and 4.0 (5th and 95th percentiles). Thus, most V4 neurons are not color-time separable. In the time separability analysis, a POP is considered time separable if its effective rank is below a value of 1.5. An effective rank of 1 would indicate perfect separability. However, it is likely that few neurons are perfectly separable, and so it is unrealistic to expect neurons to have an effective rank of 1. An effective rank of 2 indicates a more complex relationship between time and the other dimension of interest. Therefore, the cutoff of 1.5 was selected because it is a low value between 1 and 2. This cutoff is somewhat arbitrary. Therefore, we also performed a permutation test to determine if the effective rank differed significantly from a null distribution. We used the effective rank of the original pixel space of the POP to examine space-time separability. Across the sample of 313 V4 neurons, 223 neurons (71%) have a significantly lower rank than the null distribution (p < 0.025, FDR corrected; **Figure 7b1**). Thus, over two thirds of V4 neurons are space-time separable. We used the effective rank of the spatial spectrum of the POP to examine spectrum-time separability. Across the sample of 313 V4 neurons, 309 neurons (99%) have a significantly lower rank than the null distribution (p < 0.025, FDR corrected; **Figure 7b2**). Thus, almost all V4 neurons are spectrum-time separable. We used the effective rank of the color histogram space of the POP to examine color-time separability. Across the sample of 313 V4 neurons, only 102 neurons (33%) have a significantly lower rank than the null distribution (p < 0.025, FDR corrected; **Figure 7b3**). A further 5 neurons (1.6%) have an effective rank that is significantly larger than the null distribution (p < 0.025, FDR corrected). Thus, only a minority of V4 neurons are color-time separable.

These analyses show that most V4 neurons are spectrum-time separable, whereas only about one third of V4 neurons are color-time separable, and even fewer are space-time separable. Thus, spectral tuning appears to be independent from temporal tuning, but chromatic and spatial tuning both interact with temporal tuning. This may reflect well-known differences in the temporal responses of the luminance and color pathways in the primate visual system ^98^.

## DISCUSSION

The overarching goal of this study was to construct an accurate, biologically plausible computational model of single area V4 neurons that could account for their responses under naturalistic conditions. Based on the complexity of spatial and chromatic tuning in area V4, we suspected that creating such a model would require long-duration recordings from single V4 neurons. Therefore, we developed a method for recording single V4 neurons over several weeks or months (see **Supplementary Figure S1**). Using this approach we recorded a large V4 data set of unprecedented scale, consisting of 313 isolated V4 neurons, each probed with up to 1.4 million frames of naturalistic video. To model this large data set, we developed a new computational architecture that reflects the known functions of the early and intermediate visual hierarchy, and which also leverages the power of modern DNNs. The resulting *hierarchical convolutional energy* (HCE) architecture accurately predicts responses of single V4 neurons to naturalistic videos, consistent with the hypothesis that the first three stages of the cortical visual processing hierarchy consists entirely of units like those found in area V1.

One disadvantage of the HCE architecture (or indeed any deep network architecture) is that the stacked nonlinearities make it difficult to visualize or quantify receptive field tuning parameters directly from the model. Therefore, to interpret the HCE models we synthesized the optimal pattern that, according to the model fit to each individual neuron, would be predicted to maximally drive the neuron. These predicted optimal patterns (POPs) were then used to quantify the spatial, chromatic, and temporal tuning of each neuron. Statistical analyses of these POPs confirmed prior reports of shape and color tuning in area V4, and also revealed several novel tuning properties in this intermediate visual area.

### The hierarchical convolutional energy model

One of the most important contributions of this study is the development of the HCE architecture for area V4. The HCE architecture is a modular deep network that implements a hierarchy of biologically plausible computational mechanisms based on those thought to exist in visual area V1 (see **Figure 2**). To facilitate fitting models to large neurophysiology data sets we also leverage the power of modern GPU-based deep learning algorithms. The HCE architecture produces excellent predictions of V4 neuronal responses, even under naturalistic conditions and at short time scales.

The HCE architecture reflects an important hypothesis about the early stages of processing in the primate visual cortex. Specifically, it assumes that areas V2 and V4 each apply spatial filtering operations on their inputs that are analogous to the spatial filtering that is found in V1. In other words, the model assumes that the visual hierarchy, at least through area V4, is implemented as a cascade of self-similar nonlinear filtering and pooling operations like those found in area V1. This scheme is particularly elegant because it only requires computational mechanisms that are already known to exist in area V1 to account for the complex shape selectivity observed in area V4. The success of the HCE architecture implies that it might also be useful for modeling other intermediate visual areas, such as area MT. In fact, the modular nature of the HCE framework suggests it could be extended to model downstream higher-order visual areas located within inferior temporal cortex.

Previous models of V4 tuning, such as those based on curvature angle ^12^, spatial frequency ^56^, and color ^47^, were limited in scope and unable to account for responses under naturalistic conditions. Recent models employ a modern ADNN framework ^99,100^, often using transfer learning due to insufficient neurophysiological data for end-to-end training ^60–64,76,101^. These models can predict responses to flashed images but not to movies. Conversely, HMAX-inspired models, such as those by Cadieu ^57^, Mehrani and Tsotsos ^59^, and Popovkina and colleagues ^58^, use biologically-inspired architectures. However, the HCE architecture differs fundamentally from those models because the HCE architecture hypothesizes that V4 receptive field properties are a direct consequence of hierarchical stacking and mixing of simple and complex cell basis functions. Furthermore, while the earlier HMAX-inspired models were only designed to explain V4 responses to simple parametric shapes, the HCE architecture was designed to predict V4 responses to full color naturalistic videos.

On the practical side, the HCE model has relatively few parameters, especially as compared to recent ADDNs developed in the computer vision community. For example, while the VGG model used here requires 14,715,216 parameters, the HCE model requires only 20,620 free parameters. The relatively low parameter count of the HCE model enables end-to-end training directly on neural data, something that cannot be achieved with conventional DNN models.

### Spatial tuning in area V4

Our results show that V4 neurons have complex and diverse patterns of spatial selectivity. Many of the properties that we observe are consistent with previous reports of spatial tuning in V4; others are consistent with prior theoretical predictions that had not been confirmed experimentally; and some aspects of spatial tuning observed here have not been reported previously. Our results suggest that spatial tuning in V4 is not well described by any simplistic model that assumes that these neurons fall into distinct, spatially-selective categories or clusters. Instead, V4 spatial selectivity appears to reflect a continuous distribution of spatial selectivity properties across a complex space spanning both shape and texture. This diverse spatial tuning can be encapsulated using several metrics, such as the POP spatial spread, the spectral tuning, the entropy of the spectral tuning, or the inhomogeneity of spectral tuning over the receptive field.

One key aspect of spatial representation in area V4 concerns curvature. Previous studies of V4 have shown that these neurons are selective for concentric and radial gratings of different curvatures ^10,11^; for curved 2D contour elements ^13,14,34^; and for 3D curvature denoted by stereo ^26^ or by shape from shading ^27,102^. Furthermore, lesions to human V4 impair curvature discrimination ^19^. In light of these prior studies, it is reassuring that our results confirm that curvature selectivity is one of the most important aspects of spatial tuning of area V4 neurons under naturalistic conditions. That said, V4 neurons should not be viewed as simple curved edge detectors. Prior studies have shown that V4 neurons are also selective for non-Cartesian hyperbolic gratings ^10,11^, intersections ^103^, or acute angles ^12^. Our results also confirm these earlier observations. Finally, several prior studies of V4 have reported that one of the most important dimensions that differentiate these neurons is their selectivity for contours versus textures ^10,11,61,104^. This important dimension of spatial selectivity is also manifest in our data.

The success of the HCE architecture suggests that selectivity for curvature is at least partly a consequence of the log-polar transform implemented in area V1. As first noted by Motter ^55^, receptive fields of all visual pathway neurons downstream from area V1 inherit visual field sampling biases due to the cortical magnification present in the primary visual cortex. (The log polar representation in V1 itself appears to be driven largely by the gradient of photoreceptor density across retinal eccentricity.) Any neuron downstream from V1 that samples from the log polar map of V1 will have spatial receptive fields that are oblong in retinotopic coordinates. Although this property was highlighted in Motter’s original publication ^55^, the way that the log polar map in V1 induces curvature selectivity in V4 was not proposed in that publication nor by any previous studies of spatial selectivity in V4 of which we are aware. In sum, the complex spatial tuning found in area V4 neurons appears to be a consequence of both the log polar transform implemented in V1, and of the hierarchical stacked arrangement of convolutional energy filters implemented across visual areas V1, V2 and V4.

### Chromatic tuning in area V4

Given the longstanding interest in color representation in area V4, another important goal of this study was to create a biologically plausible computational model that could account for color responses of V4 neurons ^41,42,49^. To this end, we designed the HCE model so that it could account for responses to chromatic stimuli. (Note that the current HCE model encodes color in terms of the red, green, and blue channels of the monitor, and it does not aim to model color in terms of known neurophysiological color channels in the early and intermediate visual systems. However, it would be straightforward to modify the model to represent color in a more biologically plausible form.) To understand chromatic tuning in the HCE models fit to our sample of V4 neurons, the POP for each neuron was projected into a low-dimensional chromatic embedding designed to reveal different aspects of chromatic selectivity. A principal component analysis was then used to summarize the dimensions that best describe the variation in chromatic tuning across our sample of V4 neurons.

Our results show that many V4 neurons are selective for color. About one third of the neurons are selective for red-cyan contrast, about one quarter are selective for blue-yellow contrast, and about one sixth are selective for green-magenta contrast. Chromatic embedding analysis reveals that three principal components account collectively for about half of the variance in chromatic tuning across our V4 sample. These PCs discriminate tuning for black and white versus blue and yellow, blue and yellow versus green and magenta, and blue and yellow versus red and cyan. Thus, the PCs of the chromatic embedding distinguish between corners of the three dimensional color cube. This confirms many prior reports showing that color-selective V4 neurons are broadly distributed across color space, with a predominance of black-white selectivity ^47,48,91^.

Shape and color information are represented in partly segregated processing streams in areas V1 and V2 ^105^, and area V4 also appears to contain specialized structures (’color globs’) that contain a high proportion of color-selective neurons ^48^. However, chrominance and luminance information are highly intermixed in many of the V4 neurons in our sample. Projections onto chrominance and luminance components of the POPs show that the majority of V4 neurons are selective for both chrominance and luminance. Comparison of luminance and chrominance spectral entropy shows that color-selectivity is biased toward low spatial frequencies, while luminance-selectivity is biased toward higher spatial frequencies. This is consistent with a large body of evidence showing that the primate color system has lower spatial fidelity than the luminance system ^106^.

### Temporal tuning in area V4

Prior studies have paid relatively little attention to temporal response properties of V4 neurons. Most studies have used static stimuli shown for several hundred milliseconds, and they only sought to predict responses at those long timescales. However, natural stimuli are highly dynamic, and they evoke complex responses from neurons across the visual pathway that cannot be described by merely integrating response rates over long time windows^70,78,95^

To address this issue, our study used naturalistic videos as stimuli, and developed the HCE architecture to model the temporal responses of V4 neurons. In order to accurately predict fast temporal dynamics, the HCE architecture operates at the 60 Hz video sampling rate of the stimuli (16.7 ms resolution). This enables the model to predict responses at a fine time scale. However, because neuronal responses at short timescales are dominated by sub-Poisson noise ^82,83^, the noise ceiling and prediction accuracy of the HCE model is somewhat lower than has been reported by other models operating at much longer timescales of hundreds of milliseconds. We recovered temporal tuning profiles by analysis of the POPs generated from the HCE model fit to each V4 neuron. We find that the temporal responses of these neurons can be explained by three principal components. The first PC separates V4 neurons tuned for relatively sustained patterns from those tuned for relatively more transient patterns. The second PC separates neurons tuned to transient patterns based on their peak’s temporal position. The third PC is similar to the first PC, separating neurons tuned to sustained patterns from those tuned to transient patterns. The temporal performance is particularly notable given that the HCE model operates at the frame rate of the video stimuli. This fine-scale temporal information is missing from most other V4 models that only predict responses on longer timescales of hundreds of milliseconds.

### Space-color-time separability

Most prior neurophysiology studies of area V4 have focused on either space, time or color, but did not examine all three variables simultaneously. However, natural visual stimuli vary continuously in space, color and time. If V4 neurons are not space-color-time separable, then the conclusions of studies that examined only one of these three domains may not generalize to the real world. Therefore, to examine this issue we evaluated space-color-time separability across our V4 sample. We find that most V4 neurons are not space-time separable, but that they are spectrum-time separable. We also find that most V4 neurons are not color-time separable, in keeping with prior reports ^54^.

It is well known that V4 neurons with overlapping receptive fields can be tuned for a variety of different spatial frequencies ^7^, so it is not surprising that much of the differences in spatial tuning among V4 neurons can be attributed to variability in spatial frequency selectivity. Because the color system in primates is biased toward lower spatial frequencies compared to the luminance system ^93,94^, color selectivity in V4 is associated with relatively lower spatial frequency selectivity. However, V4 receptive fields are complicated and they can include both chromatic and achromatic elements. Consequently, single V4 neurons that are tuned for low frequency color can also be tuned simultaneously for high frequency luminance changes (see for example **Figures 5c-d**). This finding is in line with prior work suggesting that spatial and color tuning are largely separable in V4 ^107–110^.

### The role of area V4 in visual representation

Taken together, our results indicate that area V4 forms a very high-dimensional, intermediate representation of space, time and color. This complex representation suggests that stimulus selectivity in area V4 reflects image segmentation and grouping operations that are a critical aspect of intermediate vision ^3,11^. These operations support mapping from the simple but very general representation found in primary visual cortex into category-based representations found in inferior temporal cortex ^5^. Our data explain previous observations regarding shape, curvature, texture and color tuning in V4, but they make clear that shape tuning in V4 is much more complicated than has been suggested in prior studies.

One theory about area V4 is that it plays a critical role in breaking camouflage and in forming a cue-invariant representation of shapes and their borders ^3,49,111,112^. Our results are generally consistent with this view. For example, several of the POPs appear to be akin to amplitude-modulated filters that contain multiple, divergent frequency- and orientation-selective subunits. Amplitude modulation is a well known mechanism for breaking camouflage ^113,114^, and one long-standing proposal for how the visual system might break camouflage is to implement a stack of amplitude modulated filters ^113,114^. Thus, area V4 may play a role in helping to segment scenes under naturalistic viewing conditions.

Although area V4 neurons appear to provide a robust, cue-invariant representation of borders, they do not appear to represent shapes segmented from the background. Indeed, many of the V4 neurons in our sample are selective for complex patterns that cross object boundaries (see **Figure 4d**). This suggests that any explicit representation of object shape happens at more central stages of visual processing downstream of area V4. If so then the representation of shape in area V4 may be more closely related to shape representation in areas V1 and V2, where a minority of neurons represent illusory contours ^115^ and border ownership ^116^, than it is to later stages of visual processing, where neurons appear to represent segmented objects ^117,118^.

### Relationship between representation in area V4 and natural image statistics

Natural scenes have complex, hierarchical statistical properties that reflect the physical structure of the world, and the way that images of the world are projected onto the retina. The first-order statistics of the world describe the spatial distribution of luminance and chrominance, while the second-order statistics describe the spatial distribution of edge segments, texture patches and local motion ^119,120^. Higher-order statistics describe more complex image properties ^111^. While both first- and second-order stimulus statistics are well understood for natural images, little is known about the higher order statistical properties.

The primate visual system has evolved to exploit the statistical regularities in natural images to make inferences about the structure of the world from sparse image data ^66,67,119,120^. This process takes place through a hierarchical series of processing stages. Relatively early stages of processing such as the lateral geniculate nucleus (LGN) and area V1 represent first-order (e.g., luminance) or second order (e.g. local edges) statistics ^70,121,122^. Higher-order areas in the inferior temporal cortex represent complex shapes in a relatively cue-invariant manner ^123^. There is an interesting analogy between the hierarchical organization of the primate visual system and the hierarchical organization of ADNNs trained on image classification tasks ^68,84^. In both cases relatively earlier stages of processing represent relatively simple statistical properties, and relatively later areas represent higher-order statistical properties. This close correspondence makes sense given that both hierarchically-organized systems have evolved (or learned) to represent the hierarchically-organized statistical structure of natural scenes.

The data presented here suggest that area V4 represents the intermediate statistical structure of natural scenes. The representation is much more complicated than that found in area V1, and it is more difficult to understand than the shape-selective representations found in higher-order inferior temporal areas. Given that area V4 represents the intermediate statistical properties of natural images, it is counterintuitive to find that V4 receptive fields appear to be more complicated than those of neurons located downstream of V4. However, an analogous finding has been reported in the artificial neural network literature. DNNs trained to solve visual categorization tasks show that intermediate layers are usually more complicated and higher-dimensional than either the earlier input layers or the later output layers of the network ^68,84,124^. This does not appear to be some simple artifact of training, since DNN models that are designed explicitly to reduce overparameterization still require high-dimensional intermediate layers ^125^. Thus, complex intermediate representations may be a fundamental requirement of any hierarchically-organized system that represents the hierarchical statistical structure of natural scenes.

The complex amplitude-modulated filtering properties of many V4 neurons provides further evidence that V4 represents intermediate statistical structure. Karklin and Lewicki ^111^ developed a model that learned second-order independent components of natural scenes from natural images. Inspection of the learned filters revealed that many of them were selective for complex amplitude-modulated patterns that mixed different orientations. The similarity of the filters predicted by Karklin and Lewicki ^111^ and the POPs produced in our study is remarkable, and it suggests that area V4 represents the higher-order independent components of natural scenes.

### Potential limitations of this study

The study reported here utilized a V4 neurophysiology data set of unprecedented size, and a sophisticated yet biologically plausible computational model. The model produces excellent predictions of V4 responses under naturalistic conditions (i.e. using naturalistic videos) and at very short time scales (i.e., at the refresh rate of the monitor). Furthermore, the model generalizes to novel videos that were not used to fit the model. This strong model can provide many novel insights about visual representation in area V4. That said, like all experiments our study is still limited in several respects. First of all, although the visual stimuli used here were naturalistic, they were not natural. The videos were drawn from commercial sources, and so were biased by decisions made by the videographers during filming ^126,127^. And although the videos were displayed on a calibrated plasma monitor, the range of luminance and the color gamut of such monitors is compressed relative to luminance and color in the real world. These factors had only a minor effect on our results. Any biases due to videography were likely washed out by the enormous size of the stimulus set, and in any case area V4 is relatively insensitive to absolute luminance ^49,91^. We believe that the most likely source of display-related bias in our study is in the domain of color: although the monitor was professionally color calibrated, no quantitative colorimetry was performed.

Our recordings were made during steady fixation. Our experiments did not include natural eye movements, an important component of the natural visual behavior of primates ^128,129^. There is evidence that eye movements change response properties of V4 neurons, through out-of-band saccadic stimulation and the modulatory effects of efference copy ^128^. However, the stimulus-driven component of saccadic modulation is likely preserved in our data, because scene cuts in the videos used here affect V4 responses similarly to saccades. Furthermore, the allocation of visual selective attention likely differs between fixation and free viewing ^129^, but efference copy and attentional effects in V4 are known to be modest, accounting for about 15% of the variance in V4 responses ^130–132^. Future work should aim to expand upon the fixation condition examined here to characterize and model V4 neurons under even more naturalistic viewing conditions.

The procedure for generating the POPs is stochastic. This means that each of the POPs shown in this paper is just one sample from a family of patterns that are predicted to elicit the largest response from each neuron. To ensure that the POP was a representative sample from this family, we examined the distribution of POPs generated for each neuron from different initial conditions (see **Supplementary Figure S4**). This procedure confirmed that the POPs shown here indeed robustly represent the patterns for which each V4 neuron is tuned.

The log-polar transform used in the HCE architecture was implemented explicitly and so it was not included in the model as a layer optimized during training. If some V4 neurons don’t follow the sampling pattern described by Motter ^55^ then this implementation may be suboptimal. In theory a learnable convolutional layer with filters of increasing receptive field size could have been implemented as an additional layer in the HCE architecture, but this would have increased computational demands so it was not implemented.

Several neurophysiology laboratories have implemented a public scoring system where scientists can submit their neural models to be evaluated against neurophysiology data ^133^. (Our laboratory developed the first tool of this kind, the Neural Prediction Challenge, which we supported from 2005 to 2015.) However, these efforts are too limited to support evaluatuon of the HCE model. The visual neuroscience experiments underpinning current data challenges such as the Brain-Score project ^133^ used static stimuli, and they only aim to predict responses over a timescale of hundreds of milliseconds. In contrast, the HCE model takes in video and predicts at a resolution of 60 Hz. Thus, the current Brain-Score framework cannot account evaluate a model such as the HCE architecture.

### Conclusions

This study verifies a new computational architecture called the *hierarchical convolutional energy* (HCE) model that can account for responses of single V4 neurons (and so the neuronal population) during stimulation with full color nature videos. The HCE architecture reflects a biologically plausible hypothesis about visual processing in the primate ventral processing stream: that areas V2 and V4 each apply spatial filtering operations on their inputs that are analogous to the spatial filtering operations that occur in area V1. By stacking these operations in a hierarchical sequence, areas V2 and V4 can represent explicitly more complex aspects of shape than the elementary features represented in V1. Thus, this elegant scheme accounts for V4 responses using only computational mechanisms that are already known to exist in area V1. If computations within area V4 simply reflect a stack of V1-like computational modules, then this begs the question of how far this framework could be pushed in the visual processing hierarchy. Future work should focus on extending the HCE architecture, and apply it to other intermediate visual areas such as MT, or to visual areas further downstream in the visual processing pathway such as posterior inferior temporal cortex. Note that the modeling effort reported here would have failed had we collected a smaller dataset more typical of V4 studies. We found that model performance increased with increasing training set size. In fact, even at a size of 1 million training data points the model performance did not appear to converge. Thus, this study illustrates the necessity of collecting large, high-temporal resolution neurophysiology dataset in the big-data regime of the modern day.

### Online data viewer

The information provided in this manuscript summarizes general tuning properties across our large V4 sample. However, there is substantial variability in spatial, chromatic and temporal tuning across individual V4 neurons that cannot be easily captured by summary statistics. Therefore, to enable readers to explore the data more fully we have also created an interactive web viewer that supports exploration of the POPs and the similarity metrics used in this paper. The viewer is hosted online at <https://gallantlab.org/viewer-oliver-2025/>. The web viewer consists of two separate interactive applications. The first provides a means to view the distribution of all of the V4 neurons in our sample in any 2D projection used throughout this study, displaying the POPs, the spatial spectra, the color histograms, or the temporal response profiles. The limits of the viewer can be adjusted to select specific regions within the 2D distribution as desired. The second interactive application provides a means to visualize the tuning properties of individual V4 neurons in more depth. The POP estimated for each neuron is displayed across all 14 temporal lags, along with the spatial spectrum, the color histogram, and the temporal response profile. This viewer also displays the twenty V4 neurons that are most similar to the selected neuron. To facilitate exploration of population coding properties, the similarity metric can be controlled by the user.

## METHODS

All animal procedures were approved by the Animal Care and Use Committees at the University of California, Berkeley, and met or exceeded all National Institutes of Health and U.S. Department of Agriculture regulations.

### Visual stimulation and task

Stimuli were displayed in a dimly lit room on a professionally calibrated 60” Panasonic V300 plasma television positioned 58 cm from the eye, covering 98 degrees of visual angle horizontally. The plasma television provided a large field of view while maintaining a pixel response-time equivalent to a cathode-ray tube display. At the point of fixation, each pixel subtended 0.054° of visual angle. This spatial resolution matched that used in prior recordings from foveal V1 neurons ^134^.

Many prior studies of area V4 have used simple parametric stimuli. These stimuli efficiently sample a stimulus subspace of particular interest. However, because V4 neurons are nonlinear, models based on such recordings do not predict neuronal responses outside the tested stimulus space, and they do not accurately predict responses under naturalistic conditions ^95^. In contrast, dynamic, full-color naturalistic movies evoke responses that reflect nonlinear mechanisms that are present during natural vision. Therefore, neuronal models created using these stimuli generalize well across a broad range of natural and artificial stimuli ^70^, and they provide the most accurate predictions of responses to naturalistic stimuli ^95,135,136^. For this reason, the experiments described here used naturalistic color natural movies taken from the nature documentaries “Planet Earth” and “Life” ^126,127^.

The movies were displayed while animals fixated steadily at a single point. To simulate natural stimulation that occurs as a result of saccadic eye movements, the movies were spliced into short clips that changed randomly every ∼300 ms ^70,128,131^. Because high-speed motion is relatively rare in natural movies, it is inefficient to use natural movies directly to estimate the high temporal frequency receptive field components ^70^. Therefore, to boost the temporal frequency content of the natural movies and increase estimation efficiency, all movies were displayed at 60 Hz rather than their native frame rate of 24 Hz. An earlier study showed that this manipulation does not bias the estimated spatiotemporal tuning of neurons of the middle temporal visual area (MT) ^70^. Because V4 neurons have lower temporal tuning than MT neurons ^36,131^, this approach likely did not bias our estimates of V4 receptive fields.

During all recordings, animals performed a simple fixation task for liquid reward. During each trial, animals maintained fixation on a small red square with a black outline. When full-field natural movies were presented, the square was superimposed on the movie. Otherwise, the square was presented on a gray background. Trials lasted 3-5 s. Animals received a liquid reward if they maintained fixation within a radius of 0.5° during this period. To encourage precise fixation, larger rewards were given for maintaining fixation within a radius of 0.3°. Eye position was monitored using a 1000 Hz video eye tracker (Eyelink II, SR Research, Ontario, Canada) and the standard deviation of fixational eye movements was typically about 0.1° ^137^.

### Electrophysiological recording

Extracellular recordings were made from area V4 in two adult male rhesus macaques (Macaca mulatta). V4 was located by stereotactic coordinates and verified by visualization of the lunate sulcus. Recordings were made from a 96-channel Utah microelectrode array ^138^ placed chronically in one hemisphere of each animal (left hemisphere in one animal, and right hemisphere in the other). Signals were amplified and recorded at 24.4 kHz (Tucker-Davis Technologies).

Spike sorting was performed offline using a custom semi-automated clustering algorithm. Because the tines of the Utah array are each 400 micrometers apart, it is safe to assume that no single neuron was simultaneously recorded on more than one electrode. Therefore, the spike-sorting algorithm was applied separately to each one of the 96 electrodes and for each day of recording. Voltage traces were first band-pass filtered at [500, 6000] Hz and thresholded to extract spike waveforms. The spikes were then peak-aligned and windowed with a skewed Gaussian to retain the spike shapes while suppressing noise both before and after the spike. (Using a skewed Gaussian helps ensure that the window decays softly away from the spike amplitude maximum, while not cutting off the longer part of the waveform after the peak.) The spike waveforms were then projected onto a basis of Daubechies-4 wavelets. The Kolmogorov–Smirnov test was used to find the wavelet projections with the least normal distributions, and the subspace defined by the ten projections with the least normal distributions were used for clustering. Clustering was based on a variational Bayesian Gaussian mixture model that uses a Dirichlet prior to determine the best number of clusters. Since the results of these clustering algorithms depend on initialization, the procedure was repeated 40 times to obtain a distribution of cluster centers. The cluster centers that were reliable across different initializations were averaged and then used to seed a final clustering. To ensure that each neuron was assigned to only one cluster, the algorithm was tuned to favor relatively more clusters than were required to fit the data (i.e., up to 6 clusters per electrode waveform). Finally, spikes were binned to the 60 Hz frame rate of the stimuli. Prior to binning, spikes were convolved with a Lanczos temporal window (consisting only of a positive center lobe) of size 16.63 milliseconds to minimize uncertainty around bin edges.

### Single neuron identification across days by means of functional fingerprinting

The Utah array is a chronic implant and the electrodes are fixed relative to the recording array. Therefore, it is reasonable to assume that some subset of the neurons recorded on one day might still be isolated on subsequent days. However, because small shifts of the brain relative to the fixed recording array can have dramatic effects on the spike waveforms, these neurons cannot be tracked based on their spike waveform alone. Therefore, to aggregate spikes recorded across multiple days, we took advantage of the fact that different V4 neurons have complicated and unique spatial, chromatic and temporal selectivity ^7,11,36,48,50^. Thus, a complex naturalistic video will evoke a pattern of responses from each V4 neuron that, on average, is easily distinguished from those from a different V4 neuron, and the peri-stimulus time histogram evoked by a naturalistic movie can serve as a functional fingerprint for a specific neuron.

We designed a simple procedure to estimate a functional fingerprint for every neuron recorded and sorted on each day of recording. These functional fingerprints were used to track individual neurons across days. First, a 7.5-second (450 frame, 60 Hz) fingerprint stimulus was created, consisting of a large variety of natural images, movie clips, 1/f noise and other complex patterns. This movie was the same size as the aforementioned natural movies. The fingerprint stimulus was then shown at the beginning of each recording session while the animal performed a simple fixation task for a liquid reward. The fingerprint stimulus was repeated 10 times before proceeding to the main experiment. The fingerprint stimulus was shown at the same offset with regard to fixation as the aforementioned natural movies.

We found that each V4 neuron produced an idiosyncratic pattern of responses to the fingerprint stimulus, and that these responses were consistent across recording sessions. To ensure that neurons were correctly identified across recording sessions, they were only matched across days if the correlation between their fingerprints was unambiguous (typically requiring a correlation of 0.7 or higher, adjusted as needed for neurons with sparse activity). Fingerprints obtained on each successive day were treated the same way, enabling us to track single neurons over many days or weeks. In some cases we recorded from one V4 neuron for over 60 days. Using this procedure, a total of 194 unique neurons were identified in the first animal, and 132 neurons in the second animal.

### The hierarchical convolutional energy (HCE) model

Computational models of area V4 neurons that have been proposed in previous studies suffer from two main limitations. First, many of these models fail to predict V4 neuronal responses at short time scales and under naturalistic conditions ^12,56,57,60,62,64^. Any plausible model of area V4 must, first and foremost, accurately predict responses under naturalistic conditions ^95^. Second, previous models were biologically implausible. Some of these models were based on surface features, such as curved line segments ^12^ or spectral patterns ^56^. These abstract models suggest no obvious implementation. Other models are based on ideas drawn from the artificial neural network literature, such as the HMAX framework ^57^ or pre-trained DNNs ^61,62^. Although these models are loosely inspired by neurobiology, they do not conform to the known structure of the primate visual system and they cannot easily be implemented in known neural circuitry.

To overcome both of these limitations, we sought to develop a new computational model that could predict V4 responses under naturalistic conditions, and which could be implemented directly in known neural circuitry. Because the brain often reuses established circuits to perform new computations ^139^, we hypothesized that areas V1, V2 and V4 might implement similar sorts of computational operations. If true then this would dramatically simplify the process of creating a computational model of V4, because the computational principles and neural circuitry in area V1 are well understood (at least in comparison to area V4).

The resulting Hierarchical Convolutional Energy (HCE) architecture is designed to account for the hierarchical processing that occurs across areas V1, V2 and V4 (see **Figure 2**). The HCE architecture consists of a stack of computational subunits corresponding to simple- and complex-cells found in V1. In essence, the HCE architecture assumes that areas V2 and V4 both perform the same types of operations on their inputs as those that are performed in V1. However, by stacking these operations in sequence, areas V2 and V4 explicitly represent relatively more complex aspects of shape than are represented explicitly in V1.

Only ∼20k free parameters are required to fit the HCE architecture to data recorded from an area V4 neuron. This is a tiny fraction of the free parameters that would be required to fit a conventional DNN end-to-end. Even a purely linear model like the spike-triggered-average would have 64*64*3*14 = 172k free parameters. Thus, the HCE architecture provides an efficient means to fit a separate model end-to-end on the data recorded from each individual V4 neuron.

The fitting procedure is divided into three steps. First, the stimulus videos are cropped around the neuron receptive field. Second, the HCE model is initialized to random values. Third, the free parameters of the model are learned via stochastic gradient descent to fit the neuronal data. These three steps are described in more detail in the three following sections.

### Receptive-field estimation and log-polar transform

The first step to fit the HCE model was to estimate the receptive field of each V4 neuron. Because the display used for our experiments was very large (96 x 64 deg visual angle), each V4 receptive field covered only a small portion of the display (4 to 12 deg). Therefore, before modeling any neuron each frame was cropped to encompass the estimated classical and non-classical receptive fields of that cell. Because V4 neurons receive their inputs from the log-polar retinotopic map in V1, V4 receptive fields are somewhat elongated and fan-shaped ^55^ see **Figure 1c**). Therefore, before estimating V4 receptive fields, all video frames were log-polar-transformed with respect to the fixation point (see **Figure 1b**).

The log-polar transform resamples image space using the coordinates:

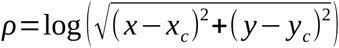

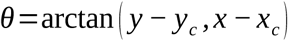

Here *ρ* is the logarithm of the distance from a point to the fixation center (*x_c_ , y_c_*), *θ* is the angle, and (*x , y*) is the Cartesian coordinate of the point that is remapped to (*ρ , θ*) (see **Figure 1b**). This transformation oversamples foveated pixels and undersamples increasingly eccentric pixels. (Note that cortical magnification could be implemented in other ways. For example, rather than log-polar-transforming the stimulus, one could build a convolutional model with radially increasing receptive field size in the first computational layer. However, this model would be inefficient to fit because it would not be able to use the heavily optimized small-filter convolution operations available in modern GPUs.)

Next the stimulus was cropped to encompass the classical and non-classical receptive field of each neuron. Different methods were used to do this for the two animals. For the first animal, the receptive field location of each neuron was estimated using an independent data set collected specifically for this purpose. Estimation stimuli consisted of synthetic stimuli such as Cartesian and non-Cartesian gratings ^10,11^ and parametric shapes ^12^. These synthetic stimuli were flashed for 83 ms at random locations while the animal fixated. The receptive field of each neuron was estimated by calculating the spike-triggered average of responses to the ensemble of synthetic stimuli. The spike time series was temporally lagged to account for the neural response delay time relative to the movie frames, and the spatial receptive field was estimated on the temporal delay with the largest spike-triggered average response.

Synthetic stimuli were not used during recording of many of the V4 neurons obtained in the second animal. Therefore, a different strategy was used to identify spatial receptive fields in that case. First, the stimulus movies were preprocessed using the pretrained AlexNet model ^68^, transforming each frame into abstract visual feature representations. Then, the spike-triggered average was computed across AlexNet’s feature activation maps to estimate the spatial receptive field. (This procedure was validated based on neurons from the first animal who saw both synthetic and natural stimuli; see **Figure S2**.) The fourth convolutional layer of AlexNet provided receptive field estimates most consistent with manually mapped receptive fields and so this was used to estimate receptive fields for each V4 neuron in the second animal. The receptive field center was taken as the center of mass of the region highlighted by the spike-triggered average. A 160×160 pixel region around the center of mass was then extracted and downsampled to 64×64 pixels. (We used the same large crop of 160×160 pixels across all neurons because the projection of V4 receptive fields onto the log-polar axes normalizes their receptive field sizes, and because we did not want to include stimulus features that were far outside the receptive field of the neuron; see **Figure 1b**.) This final 64×64 pixel image was used as input to the HCE model (see below). As with the first animal, the spike time series was temporally lagged to account for the neural response delay time relative to the movie frames, and the spatial receptive field was estimated on the temporal delay with the largest spike-triggered average response.

### Core modules of the HCE architecture

The core of the HCE architecture consists of a sequence of three modules: the V1 module, the V2 module, and the V4 module. The weights of these modules are optimized separately and simultaneously to estimate a separate HCE model for each V4 neuron (see **Figure 2a**). The V1 module (see **Figure 2a**, orange panel) implements a collection of spatiotemporal filters like those found in V1 simple and complex cells ^74–77^. This module consists of a set of space-time-separable filters, followed by squaring and rectifier nonlinearities. The set of spatial filters comprises 8 real-valued filters (as in V1 simple cells) and 8 complex-valued filters (as in V1 complex cells), each applied to the input video sequence. All spatial filters have size 7×7 pixels and the complex-valued filters are regularized towards being quadrature pairs. Note that 8 spatial filters were used in order to maximize performance by overparameterization while ensuring that the model will fit in GPU memory, and the filter size was set at 7×7 pixels to maximize the tradeoff between computational efficiency and filter interpretability. The temporal filters consist of 15 complex-valued filters of length 14 frames. These filters are created by multiplying a set of 5 cosine and sine function pairs at certain learnable frequencies by a set of 3 learnable temporal envelopes. This yields 15 amplitude-modulated complex-valued oscillatory time series. The spatio-temporal filter outputs are squared and summed, similar to those in the canonical V1 energy model ^121^. The V1 module consists of responses that reflect all possible combinations of spatial and temporal filter pairs, for a total of 240 spatiotemporal filter outputs. To select the relevant components of the V1 module and to restrict the number of parameters that must be processed by subsequent modules, the signal output by the V1 module is reduced in dimensionality from 240 channels to 20 channels using a learnable 1×1 pixel spatial convolution layer.

The second module of the HCE architecture, the V2 module (see **Figure 2a**, red panel), also implements a collection of spatiotemporal filters like those found in V1 simple and complex cells. However, these filters are applied to the efferent signals arriving from the V1 module. Our motivation to use a V1-like representation to model V2 was two-fold. First, relatively few computational models of area V2 have been proposed ^140–142^, and those that have been proposed cannot account for V2 responses during naturalistic stimulation. Second, because the computational principles underlying processing in V1 are well understood, a V2 model based on the same principles would be both evolutionarily plausible and easily interpretable. As in the V1 module, the V2 module consists of a spatial convolution with 8 real-valued spatial filters and 8 complex-valued spatial filters, applied to each channel of the input separately. All filters are of size 7×7 pixels. This operation increases the number of channels multiplicatively from 20 channels to 320 channels. Therefore, to select the relevant components of the V2 representation and to restrict the number of parameters that must be processed by subsequent modules, a learnable 1×1 pixel spatial convolution layer is used to reduce the signal output by the V2 module in dimensionality to 20 channels. With V1 temporal filtering reducing time to a singleton, the V2 and downstream modules contain no further temporal filters, proceeding in lockstep with V1 temporal processing.

The third module of the HCE architecture, the V4 module (**Figure 2a**, pink panel), consists of a set of 10 spatial receptive field masks of size 12×12 applied to the efferent signals arriving from the V2 module. Each spatial receptive-field mask produces 10 features for each input channel, totalling 200 features. Then, two separate fully-connected linear layers reduce the dimensionality from 200 to 10 features and then from 10 features to one signal corresponding to the spike rate prediction. The first linear layer combines the receptive-field mask features and passes them through a rectified linear unit activation. This acts as a simple/complex cell operator on the V2 module inputs. The final layer uses a parametric softplus activation, serving as a physiologically plausible spiking nonlinearity ^143^. **Figure 2c** summarizes the full HCE architecture and the number of parameters required for each module.

In order to regularize the HCE model during training, a custom dropout layer is added before each dimensionality reduction layer: one in the V1 module, one in the V2 module, and two in the V4 module. The most biologically plausible choice for a dropout layer would be to add Poisson noise. However, for signals of small amplitude, Poisson noise can push the signals to zero most of the time, blocking the gradient propagation during model training. To avoid this issue, a Gaussian approximation of Poisson noise is used. In the original Gaussian dropout layer ^144^, multiplicative Gaussian noise *_N_* (_1, *σ*_^2^) is introduced to the signal for regularization during training, transforming the signal *a* (*t*) at time *t* into *a* (*t*) *N* (1 *, σ* ^2^)=*N* (*a* (*t*) *, a* (*t*)^2^ *σ*^2^). Because of its multiplicative nature, the standard deviation of this dropout noise scales linearly with the spike rate *a* (*t*). Thus, this dropout noise does not optimally model Poisson noise, where the standard deviation of the noise scales with the square-root of the spike rate. To better model Poisson noise, we introduce a *coupled Gaussian dropout layer*, which uses additive Gaussian noise with a standard deviation that scales with the square-root of the spike rate, transforming the signal *a* (*t*) at time *t* into 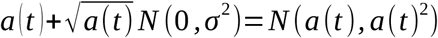. Compared to a Gaussian dropout layer, the coupled Gaussian dropout layer adds proportionally less noise when the signal is high. Indeed, if the signal increases from *a* (*t*) to 2 *a* (*t*), the standard deviation of the noise increases only by a factor 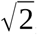, and so the signal-to-noise ratio increases by a factor 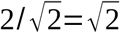. Because the noise scales with the mean of the signal, the coupled Gaussian dropout layer follows the Poisson regime and it regularizes proportionally less when the signal is high. Note that the coupled Gaussian dropout layer is only used during training in order to prevent overfitting and to increase the generalization performance of the model. For the HCE model we found coupled Gaussian dropout to be a better regularizer than standard dropout or simple Gaussian dropout, both in terms of the predictive performance of the trained models and the smoothness and interpretability of the filters learned as components of the models.

### Using the HCE architecture to model responses of individual V4 neurons

To model the responses of individual V4 neurons, the HCE model parameters were optimized by stochastic gradient descent. The goal of this procedure was to optimally predict the responses of each V4 neuron at the temporal resolution of the stimulus video (60 Hz). To predict neuron responses at a given time, the HCE model takes into account the past stimulus over a temporal window of 14 frames (233 ms). To account for the neural processing time required for the stimulus signal to reach V4, the models were fit with time delays from 2 to 16 frames (33 to 266 ms; ^145^).

The experiments presented here were designed to collect two separate data sets for each neuron: a training set for estimating model parameters and a test set for evaluating and validating model predictions. The estimation set was used to fit a separate model for each neuron by minimizing a Poisson loss between the model prediction and the measured average spike rate. The model was shown the entire estimation set in batches of 400 samples. For each batch, the gradient of the Poisson loss was calculated, and the weights of the model were updated accordingly. This process was repeated for 300 passes over the entire estimation set. The HCE model was optimized using the FTML optimizer ^146^. This optimizer was chosen because it provides stable training solutions, building upon the advantages of two leading optimizers, Adam ^147^ and RMSprop ^148^.

Video stimuli were shown to the animals multiple times, so that responses were recorded to as many as 21 stimulus repetitions for each neuron. To increase the signal-to-noise ratio, the neuron responses were averaged over these repeated stimulus presentations. As a measure of the quality of a training sample, the number of repetitions for a stimulus-response pair was provided to the HCE model as a weight on that sample. This encouraged the HCE model to prioritize learning from samples with higher the signal-to-noise ratio over samples with low signal-to-noise ratio.

Because the HCE architecture was designed iteratively over time, it was important to ensure that the architecture was not overfitting to our sample of V4 neurons. Therefore, all development of the HCE architecture was performed using data from only one of the two animals. All of the data from the second animal were held out to be used as a generalization test once the HCE architecture was finalized. After the HCE architecture was fully developed and the optimization methods for fitting the model to a neuron were well established, the data from the second animal were unmasked and the remaining neurons were modeled using the HCE architecture. A similar pattern of results was obtained for both animals, indicating that the HCE architecture was not overfit (see **Figure 3b**).

### The VGG-Features model architecture

Some previous models of area V4 have used features from a pre-trained DNN as input to a general linear model ^61,62,64,99,101,149,150^. This approach assumes that the feature maps from a pre-trained DNN can provide a dictionary of features useful for modeling real neurons. Therefore, to help validate the HCE model presented here, we compared it directly to an alternative model created using features that had been learned by a well-known pre-trained DNN, VGG-16 ^84^. VGG-16 is a convolutional DNN developed to perform an image categorization task ^84,151^. It has been used as a source of features in previous studies that have modeled neurons in area V1 ^152^ and V4 ^62^.

VGG-16 consists of multiple convolutional, rectifying, and pooling layers, but not all of the layers contain information useful for modeling responses of V4 neurons. Therefore, we developed a model architecture, here called the VGG Features architecture (VGG-F), in which the layer index for feature extraction is systematically adjusted as a hyperparameter. As in the HCE architecture described above, each frame of the stimulus video is first log-polar-transformed (see **Figure 2b**, blue panel). The transformed stimulus is then fed to a feature extraction module that uses the pretrained VGG-16 DNN to extract features from each frame of the stimulus video. In this module, each frame is passed through the first *L* layers of the VGG-16 network, producing feature activation maps of size *Height _L_* \**Width_L_* \**Channels _L_* (see **Figure 2b**, yellow panel). Because the original VGG-16 network was only trained on static images, this stage also includes a temporal delay *D* to map the output at time *T* from the video frame at time *T − D* (see **Figure 2b**, yellow panel). In this study the hyperparameters *L* and *D* were selected individually for each V4 neuron, using a held-out data set that was not used for training nor for testing the VGG-F model.

The final stage of the VGG-F architecture consists of a linear combination of pixels and feature channels that is optimized to predict responses of each V4 neuron in a held out data set. First, a receptive field mask of size *Height _L_* \**Width_L_* is learned and averaged in order to produce a single activation per feature channel. This mask is then fed into a fully-connected linear layer whose weights are optimized to maximize prediction accuracy (see **Figure 2b**, purple panel). The resulting transform is rank-1 separable in space and feature channels. The VGG-F model was trained using the Adam optimizer ^147^. The VGG-F model consists of learning a linear layer over a pre-trained DNN so the model converges much faster than would occur in a DNN model trained end-to-end. Therefore, the Adam and FTML optimizers perform similarly here and there is no compelling reason to prefer one over the other.

### Estimating prediction accuracy and statistical significance for the HCE and the VGG-F models

The normalized correlation coefficient (*CC_norm_*) ^153^ was used to quantify prediction accuracy of the fit HCE and VGG-F models. This metric provides a measure of prediction accuracy relative to the noise ceiling inherent in the data ^154,155^. *CC_norm_* is computed using the following procedure. First, the absolute correlation coefficient (*CC*_abs_) is computed between the model prediction (*X*) and the average (*Y*) of the repeated trials in the test set: 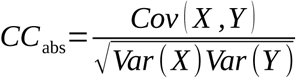. Second, the signal power (*SP*) is computed, quantifying the repeatability of the spike train (*R_n_*) evoked by the stimulus in the test set: 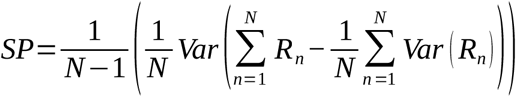. Third, the total power (*TP*) is computed, quantifying the average of the signal variance across repeats, 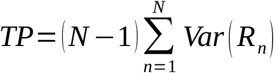. Fourth, the signal power and the total power are combined to estimate the noise ceiling (*CC_max_*), which corresponds to the maximum obtainable correlation coefficient given the repeatability of the signal: 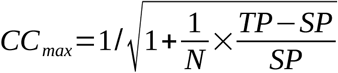. Finally, the normalized correlation coefficient (*CC_norm_*) is computed by dividing the absolute correlation coefficient by the noise ceiling: 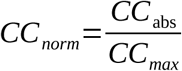.

Normalizing the performance in this way makes it possible to compare the model prediction accuracy across the entire population of V4 neurons, regardless of the signal quality of each individual neuron. Thus, this metric maintains the intuitive range of the Pearson correlation coefficient while adjusting for the signal quality of the data. Note that the *CC_norm_* and *CC_max_* could not be computed for 10 of the 326 V4 neurons in our sample because their signal power estimate was negative due to high noise. These neurons were discarded before analysis, leaving 316 neurons.

To compare the prediction accuracy of the HCE and VGG-F models, a paired-sample permutation test was used to test the hypothesis that the HCE model outperformed the VGG-F model across all recorded neurons. For each permutation, the assignment of HCE versus VGG-F was randomly permuted for each V4 neuron in the sample, and the difference in prediction accuracies was computed. This procedure was repeated 100,000 times to generate a null distribution corresponding to the null hypothesis of no difference in model prediction accuracy between the two models. Statistical significance was evaluated by determining whether the obtained difference in prediction accuracy across the sample exceeded 95% of the permuted differences in prediction accuracy, corresponding to a one-tailed paired-sample test at *α* = 0.05.

### Visualization of V4 receptive field properties

The HCE model is a hierarchical cascade of computational modules meant to reflect key processing stages of the early and intermediate visual hierarchy. While this hierarchical architecture provides accurate predictions of neuronal activity, it comes at the cost of interpretability: the receptive field properties of individual V4 neurons cannot be visualized directly by examining the coefficients of the fit HCE model. Therefore, to visualize receptive field properties of individual V4 neurons, we used a gradient-based optimization procedure to synthesize the pattern that would be predicted, based on the fit HCE model, to elicit the largest possible response from each neuron. This gradient-based visualization approach is a widely used technique when working with differentiable neural network models to model neural activity ^62,63,156^. (Note that because the HCE model contains phase-insensitive complex cells, in most cases the phase component of the recovered POPs is underdetermined.)

To initialize the stimulus optimization procedure, we first searched through the data recorded from each neuron to identify the 14-frame (233 ms) stimulus segment that elicited the strongest observed response. An iterative procedure was then used to modify this initial segment in order to maximize the response of the HCE model fit to that neuron. The loss function minimized in this iterative procedure was the negative of the predicted response of the HCE model, leading to a maximization of the predicted firing rate. At each iteration, the gradient with respect to this loss was used to iteratively update this 14-frame sequence. This procedure produced synthetic 14-frame movies that were predicted to maximally drive the responses of each V4 neuron. Here we refer to these movies as predicted optimal patterns (POPs).

To assess the stability of the POP generation procedure, the stimulus optimization was initialized using 100 different stimulus segments chosen at random with the constraint that the randomly chosen segments had elicited a wide range of different responses. This produced a sample of 100 alternative POPs for each neuron. Qualitative and quantitative analysis of these alternative POPs revealed that the variability of POPs within each neuron was small compared to the variability of POPs across neurons (see **Supplementary Figure S4**). Therefore, all subsequent population-level analyses focus on the single POP obtained by initializing with the most effective video sequence for each neuron.

To ensure that the POPs did not include any artifacts at the border of the frame, each POP was manually checked to determine if it faded smoothly to zero at the edges. This was true for 313 out of 316 of the POPs. For the three remaining neurons, the POPs contained sharp details at the borders. These sharp details were visually similar to parts of the frames used to initialize the optimization, suggesting either that the optimization procedure failed or that the receptive fields of these neurons were much larger than what was estimated. To avoid potential confounds, these three neurons were removed from subsequent analyses.

To facilitate visualization of the POP generated for each neuron, the pixel values of each POP were first scaled to the interval [-0.5, 0.5] by applying the function , where *µ* is the maximum absolute value over all pixels and channels. (For visualization only, a bias of 0.5 was added to have values in [0, 1].) Next, the inverse log-polar transform was applied to transform the POPs into Cartesian coordinates. To facilitate comparison of POPs across neurons, the inverse log-polar transform did not use the specific receptive coordinates of each neuron, but rather used a standard set of coordinates (i.e., center position and crop angles) across all neurons. This is analogous to the size normalization procedure that is commonly applied to the spatial receptive fields of visual neurons before other tuning properties are compared across neurons. (The POPs estimated for the second animal were then flipped across the vertical meridian in order to compensate for the fact that the recordings in the two animals came from different hemispheres.) Finally, the POPs were rescaled to 120×120 pixels to facilitate subsequent statistical tests.

The stimulus optimization procedure produced a tensor of size [120 pixels, 120 pixels, 14 time lags, 3 color channels] for each of the 313 V4 neurons in our sample, reflecting a spatial-chromatic-temporal pattern predicted to elicit the maximal response from the neuron. This tensor was used for analysis of the distributional statistics of spatial, chromatic, and temporal tuning properties described below. To provide a more compact visualization, some figures only display the POP frame with the strongest norm over pixels and channels.

### Characterizing the distribution of spatial, chromatic and temporal tuning properties in V4

One important goal of this study was to gain insight about the spatial, chromatic, and temporal tuning properties of the V4 neural population as a whole. In neurophysiology, a common approach for addressing this goal involves first identifying an appropriate tuning subspace, and then characterizing the distribution of neurons within that subspace. To do this, we derived several different embeddings of the distribution of POPs estimated from the HCE models fit to our V4 sample. We also developed several metrics that describe different aspects of these distributions, along with appropriate metrics that summarize the similarity between any pair of POPs within these subspaces. The following subsections describe these different embeddings and metrics.

#### Analysis of spatial tuning across the neuron population

We used a multi-metric approach to characterize the distribution of spatial tuning properties observed in the POPs across our sample of V4 neurons. Each POP was considered in both the spectral domain and the pixel domain, and several different metrics were calculated within each domain. To create a spectral embedding, each POP was first centered on its mean value over pixels, lags, and channels. Then the 2D Fourier transform was applied to each lag and channel independently, using zero-padding to obtain a 128×128 spectrum. The magnitude of each frequency was then computed and averaged over all lags and channels. The spatial spectrum was then shifted so that the low frequencies were located at the center of the frequency plane. Finally, to reduce noise and to increase robustness of the spectral embedding, the 128×128 spectra were downsampled with anti-aliasing to size 31×31. The resulting spatial spectrum captures the spatial-spectral tuning of a neuron in a 961-dimensional embedding.

The first spectral metrics were obtained by using principal component analysis (PCA) to recover the spatial and spectral dimensions that described the largest variance in spatial tuning across the sample. The spectra were first flattened into 1D vectors and then normalized so that each vector summed to one. To ensure that neurons with the best predictions under the HCE model would have more influence in this analysis, prior to PCA the spectrum of each neuron was weighted by its clipped normalized correlation score, *w*=*max* (0.01*, C C_norm_*), where the maximum operator chooses the larger of either 0.01 or the normalized correlation score. Bootstrapping was used to assess stability of the obtained principal components (PCs) ^157,158^. In the bootstrapping procedure, the PCA was recomputed on subsets of 323 neurons sampled randomly with replacement, and then the correlation coefficient was used to compare the original PCs with the bootstrapped PCs. To assess the stability of each original PC, each original PC was correlated with each bootstrapped PC, and the maximum correlation over the bootstrapped PCs was considered. For each original PC, the median value across 1000 bootstrapped samples was taken as the measure of stability. A threshold of 0.9 was used to identify stable results. For the spatial spectral embedding, the first four PCs were considered stable (median correlation of 0.97, 0.96, 0.93, 0.92, first quartile of 0.94, 0.91, 0.9, 0.84, and third quartile of 0.99, 0.98, 0.96, 0.96).

When inspecting the POPs across our sample, we noticed that the spatial spectra of some neurons were highly concentrated in few spatial frequencies, while the spectra of other neurons were spread across many different spatial frequencies. To quantify the extent of this spread across neurons, we developed a second metric that describes spatial receptive field properties in terms of spectral entropy. In this metric, the normalized spatial spectrum embedding of the POP for each neuron was considered as a probability distribution *q*, and the entropy of this distribution was computed as *H* (*q*)=−*Σ_frequencies_ q* log(*q*). According to this metric, a neuron with a spatial spectrum spread over many frequencies will have high spectral entropy, while a neuron with a spatial spectrum concentrated over a few frequencies will have low spectral entropy.

Both spatial spectrum and spectral entropy describe the global organization of the receptive field of each POP, but they do not provide information about the spatial heterogeneity of each POP. To quantify this property, we developed a third spatial receptive field metric. First, 1000 patches of size 31×31 pixels (about a fourth of the size of the POP) were extracted from each POP at each of the 14 lags. Patches were sampled uniformly with the constraint that each patch must fit completely within the fan-shaped contour defined by the inverse log-polar transform. Next, a spatial spectrum was computed on each patch independently. Zero padding was used to create a 32×32 spectral window and spectral magnitude was averaged over channels. A singular value decomposition (SVD) was then performed over the 14000 patches in order to extract the spectral components that are shared over patches. Finally, the ratios of explained variance of all components were reduced to one value by computing the effective rank ^97^ *r* =exp (*H* (*q*)), where *q* is the vector of singular values normalized to sum to one, and *H(q)* the entropy of *q*. The effective rank is a continuous measure that quantifies the number of components that predominate in the SVD. Its value is equal to *k* when the variance is equally split between *k* components, and it continuously interpolates these values when the variance is split unequally. In the case of the distribution of spatial spectra over patches, a low effective rank indicates a homogeneous POP in which the spatial spectrum is similar across patches. A high effective rank indicates a heterogenous POP in which the spatial spectrum is variable across patches.

All the metrics described above summarize different attributes based on a spectral embedding of each POP. However, the spectral domain does not capture the distribution of receptive field properties in the pixel domain. Therefore, we supplemented the three spectral metrics with a metric that we call spatial entropy. First, the norm of each pixel comprising the POP was computed as 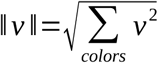, where *v ∈*[− 0.5,0 .5]^3^ is the RGB vector of one pixel. Then the norm values over pixels was considered as a probability distribution *q* normalized to sum to one, and the entropy of this distribution was computed as 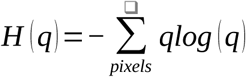. According to this measure, a neuron with a POP spread over many pixels will have high spatial entropy, whereas a neuron with a POP that is concentrated only over a few pixels will have low spatial entropy.

To estimate the similarity between POPs based on the seven metrics described above (i.e., the first four spectral principal components, the spectral entropy, the effective rank of spectra over patches, and the spatial entropy), the metrics were concatenated into vectors, and each dimension was normalized to have zero-mean and unit-variance. The similarity of the spatial receptive fields between each pair of V4 neurons in the sample was then defined as the euclidean distance between the corresponding vectors.

#### Analysis of chromatic tuning across the neuron population

We used a multi-metric approach to characterize the distribution of chromatic tuning properties found in the POPs estimated across the sample of V4 neurons. To create a chromatic embedding for each POP, a color histogram was computed for each neuron. To ensure that the embedding of color remained close to the input space of the HCE model, all color transformations were kept linear with respect to the red-green-blue (RGB) color space. To facilitate visualization when computing the color histogram, color vectors in RGB space were first shifted by subtracting 0.5. Then, color vectors were decomposed into their normalized direction on a 2D color sphere and their vector length as a 1D color intensity. The 2D color sphere was then partitioned into 100 bins of approximate equal surface area (Fibonacci lattice ^159^). The value associated with each bin corresponded to the sum of the squared length of the color vector over all pixels within the bin, 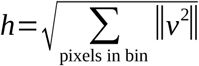. The resulting color histogram captures the chromatic tuning of a neuron into a 100-dimensional embedding.

The first color metric was obtained using a PCA procedure analogous to the one described earlier to summarize the spatial spectra. The color histograms were flattened into 1D vectors, and normalized to sum to one. These vectors were submitted to weighted PCA, where the weights were given by the clipped prediction performance of the neuron, 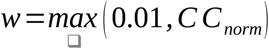. The same bootstrapping method described earlier for the spatial spectra was used to assess stability of the obtained PCs, and the first three color PCs were considered stable (median correlation of 0.94, 0.88, 0.92, first quartile of 0.86, 0.8, 0.87, and third quartile of 0.97, 0.94, 0.95).

PCA provides a data-driven approach for finding important dimensions that describe the largest variance in chromatic tuning across the sample. It may be difficult, however, to interpret these dimensions in terms of the RGB channels that encoded color information in the stimulus and which provide color channels in the HCE model. Therefore, we developed a second method that more directly recovered chromatic tuning along the RGB directions in color space. In this embedding, pixels from the POP were projected on each one of the four RGB cube diagonals: the white-black diagonal, the red-cyan diagonal, the green-magenta diagonal, and the blue-yellow diagonal. (Note that because there are only three dimensions in RGB space, these four projections are not independent.) These projections were computed using the formula, *p* =(*v^ᵀ^ D*) *D*, where *D* ∈*ℝ* ^³^ is either [1, 1, 1]^T^ (white-black), [1, −1, −1]^T^ (red-cyan), [−1, 1, −1]^T^ (green-magenta), or [−1, −1, 1]^T^ (blue-yellow), normalized to unit norm. Then the ratio of explained variance of each projection was computed as 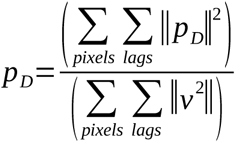

Because the HCE model architecture does not favor one color direction over another, if the pixels were sampled uniformly on the RGB cube then each diagonal would explain one third of the variance. We can investigate whether neurons are actually tuned for a color direction by inspecting the empirical distribution of pixels over all POPs. Before building this empirical distribution, pixel correlations were reduced by downsampling each POP from 120×120 to 10×10 pixels. The pixels were then shuffled over all neurons and lags before computing the explained variance of each diagonal. This process produces a null distribution that would be expected if neurons were not color-selective. Neurons whose explained variance on a given direction in RGB color space was significantly larger than this null distribution were considered to be selective for this color direction (p < 0.05, FDR corrected).

One longstanding question in V4 neurophysiology concerns whether V4 neurons are space-color separable ^49^. To address this question we therefore performed a separate analysis to quantify this property. Each POP was first decomposed into two complementary projections: the projection on the white-black diagonal, and the projection on the corresponding orthogonal plane. These two complementary projections resulted in two versions of each POP: one that reflects luminance and one that reflects chrominance. (Note that both projections are designed to be linear with respect to RGB space and invariant to permutations of the red, green, and blue channels. Thus, these two projections are not perceptually uniform.) Each version was used to compute a separate spectrum using methods described above for assessment of spatial receptive field properties. The distribution of spatial frequencies reflected in the luminance and chrominance projections was then calculated separately using the spectral entropy measure described earlier. Finally, the luminance spectral entropy and the chrominance spectral entropy were compared.

To visualize the color histograms the binned 2D color sphere was rotated so that the black-white direction would align with the vertical axis. The sphere was then flattened using a Robinson-like projection ^160^, and each bin was assigned a reference color using the closest RGB color from the bin centroid. Finally, for each color of the histogram, the reference color *_b_* _∈[ 0,1]_^³^ in each bin was faded to gray using the histogram values, 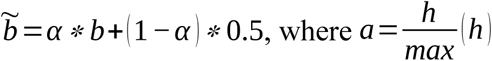.

To visualize the spectral difference of the luminance and chrominance color projections, both spectra were superimposed using a 2D colormap. In this colormap, red corresponds to high luminance spectrum values, blue corresponds to high chrominance spectrum values, white corresponds to high values in both spectra, and black corresponds to low values in both spectra.

### Analysis of temporal tuning across the neuron population

We used a multi-metric approach to characterize the distribution of temporal tuning properties found in the POPs estimated across the sample of V4 neurons. The temporal response profile of each V4 neuron was recovered from the POP estimated for that neuron. First, the norm of the POP was computed at each time lag 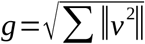, and then the norm of all 14 lags were used to form a temporal profile. The temporal profile captures the temporal tuning of a neuron into a 14-dimensional embedding.

The first temporal metrics were obtained using a PCA procedure analogous to the one described earlier to summarize spatial and chromatic tuning properties. The temporal profiles for all neurons were normalized to sum to one, and these were used in a PCA. The PCA was weighted by the clipped prediction performance for each neuron, *w* =*max* (0.01*, C C_norm_*). Stability of the principal components was assessed using bootstrapping as described above. The first three principal components of the temporal response profiles were considered stable (median correlation of 0.97, 0.99, 0.96, first quartile of 0.9, 0.98, 0.93, and third quartile of 0.98, 0.99, 0.98).

Averaging the temporal profiles of all neurons for the two animals separately revealed a small difference between the temporal response across the animals. Specifically, neurons from the first animal appeared to have a slower temporal response than neurons from the second animal. Because the 16.7 ms video frame rate is too coarse to precisely quantify this difference, the temporal response of each neuron was approximated using a Gaussian window fitted on five frame lags that were centered on the temporal profile maximum. (Similar results were obtained using a triangular or cosine window.) The Gaussian window was controlled by three parameters that specified the center, width, and height of the window. These three parameters were optimized using gradient descent and least square error minimization. The peak delay was estimated using these Gaussian window fits. Neurons with peak delays longer than 100 ms were discarded (33 neurons, 11%) and then the median peak delay was estimated across neurons separately for each animal. Statistical significance of the difference in median peak delay was assessed by means of a permutation test. In the permutation test, all neurons were randomly attributed to each animal, and the difference of median peak delay was recomputed. Repeating this procedure 10000 times led to a distribution of values corresponding to the null hypothesis that the median peak delays were identical across animals. Statistical significance was defined as the original difference of median peak delay being above the 97.5th percentile of the null distribution (two-tailed test at *α* = 0.05).

An analysis of temporal response profiles in isolation will only produce a valid result if the temporal response of a neuron is separable from other dimensions for which that neuron is selective. However, there is little evidence that V4 neurons are space-time or color-time separable. Therefore, we developed another metric that directly links space, color, and time. To compute this metric, three different embeddings were recomputed on each time lag independently. The first embedding used the original pixel embedding, the second embedding used the spatial spectra, and the third embedding used the color histograms. For each of the three embeddings, an SVD was applied over the 14 time lags and the effective rank was computed ^96^. This procedure produced a space-time effective rank, a spectrum-time effective rank, and a color-time effective rank, with values in the interval [1, 14]. According to this metric, neurons with similar embeddings over time lags will have a low effective rank, while neurons with different embeddings over time lags will have a high effective rank. A perfectly time-separable neuron (that is a neuron with a constant embedding scaled by a different intensity at each time lag) would have an effective rank equal to one.

One limitation of the effective rank metric is that it depends on the amount of total variation in the given embedding. For example, if all POPs had an almost identical color histogram, then the color-time effective rank would be trivially close to 1 for all neurons. Thus, a permutation procedure was used to determine whether the effective ranks were significantly low-rank. The effective ranks quantify self-similarity of each POP over time lags. To remove self-similarity over time lags, the POPs were permuted between neurons at each time lag. This created new permuted POPs of the form, *permuted = [neuron_a(lag_1), neuron_b(lag_2), … neuron_n(lag_14)]*. However, this permutation procedure changes the distribution of temporal profiles. For example, if all temporal profiles were pure spikes of the form [1, 0, …, 0], [0, 1, 0, …, 0], …, the permutation would create new temporal profiles such as [1, 1, 0, …, 0], or [0.5, 0.5, 0, …, 0] after renormalization. Because the effective ranks are directly dependent on the temporal profile lengths, it is important to preserve the distribution of temporal profiles in the permutation procedure. To do so, each permuted stimulus was renormalized to have a temporal profile from the distribution of existing temporal profiles. The image at each lag was renormalized with 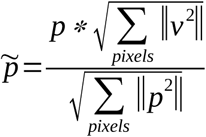. Finally, the space-time effective rank, the spectrum-time effective rank, and the color-time effective rank were computed on the permuted POPs. This resulted in three distributions of effective rank values, under the null hypothesis that there is no low-rank structure over time. Statistical significance was defined as any effective rank outside the interval defined by the 2.5 and 97.5 percentiles of the distribution (corresponding to a two-tailed test at *α* = 0.05). P-values were corrected for multiple comparisons using the FDR ^161^.

### Data analysis implementation

The data preprocessing pipeline – spike sorting, the functional fingerprinting, and the temporal downsampling – was implemented in MATLAB. The rest of the data analysis pipeline was implemented in Python, building on the scientific Python ecosystem. The HCE model was implemented using Keras ^162^ and the Theano backend ^163^. The VGG-F model was implemented using Pytorch ^164^. OpenCV ^165^, Imageio ^166^, Pillow ^167^ and H5py ^168^ were used during data preprocessing. GNU Parallel ^169^ was used in model fitting. The data analysis made use of Numpy ^170^, Matplotlib ^171^, Scipy ^172^, Scikit-learn ^173^, Scikit-image ^174^, Pandas ^175^, and Cottoncandy ^176^.

## Acknowledgments

This work was supported by the following grant awards: NIH NEI EY0169; NIH NEI EY021710; NIH NEI EY0122415; NIH EY019684; NIH B-U01EB0; NIH T32EY007043; ONR N00014-15-1-2861; ONR N00014-22-1-2217; ONR 60744755-114407-UCB; and by internal UC Berkeley funds.

## Author contributions

MO and JG conceived the experiments. MO performed neurophysiological recordings from both animals. MO wrote the spike sorting and fingerprinting software (other experimental software was developed previously as part of other projects in the lab). MO performed spike sorting and fingerprinting. MO performed stimulus pre-processing on the first animal. MW and ME performed stimulus pre-processing on the second animal. MO developed and implemented the hierarchical convolutional energy architecture, and the method for fitting the HCE architecture to V4 neurons. MW fit the HCE architecture to all V4 neurons. MW and ME implemented the VGG-F architecture, and the method for fitting the VGG-F architecture to neurons. MO and MW developed the POP optimization methods. MW performed the POP optimization for each V4 neuron. TD developed and performed all population analyses. TD developed the online data viewer. JG obtained funding, directed, and supervised all aspects of the project. The manuscript was written primarily by MW, TD, and JG, and then jointly edited by all authors.

## Data availability

Data for this study can be found at the DANDI archive (https://www.dandiarchive.org/). A digital explorer of the POP embeddings for this study can be found at <https://gallantlab.org/viewer-oliver-2025/>.

## Code availability

Code for the HCE model is available at <https://github.com/gallantlab/hce_v4>.

## SUPPLEMENTARY MATERIALS

### Tracking neurons over days with spike sorting and fingerprinting

The construction of the large dataset employed in this study necessitated multi-day neuronal recording. Thus, spiking activity was recorded over many days across 96 electrodes of a Utah array. To identify individual neurons from the recordings, a spike-sorting algorithm was applied separately to each one of the 96 electrodes and for each day of recording (see **Methods**). The spike-sorting algorithm uses the fact that spikes from the same neuron have the same waveform, thereby grouping all spikes originating from a single neuron. However, the spike-sorting approach is not suitable to track neurons across days, because small shifts of the brain relative to the Utah array can have large effects on the spike waveforms (see **Figure S1a**). Consequently, an alternative strategy was adopted for multi-day neuron tracking.

**Figure S1:**
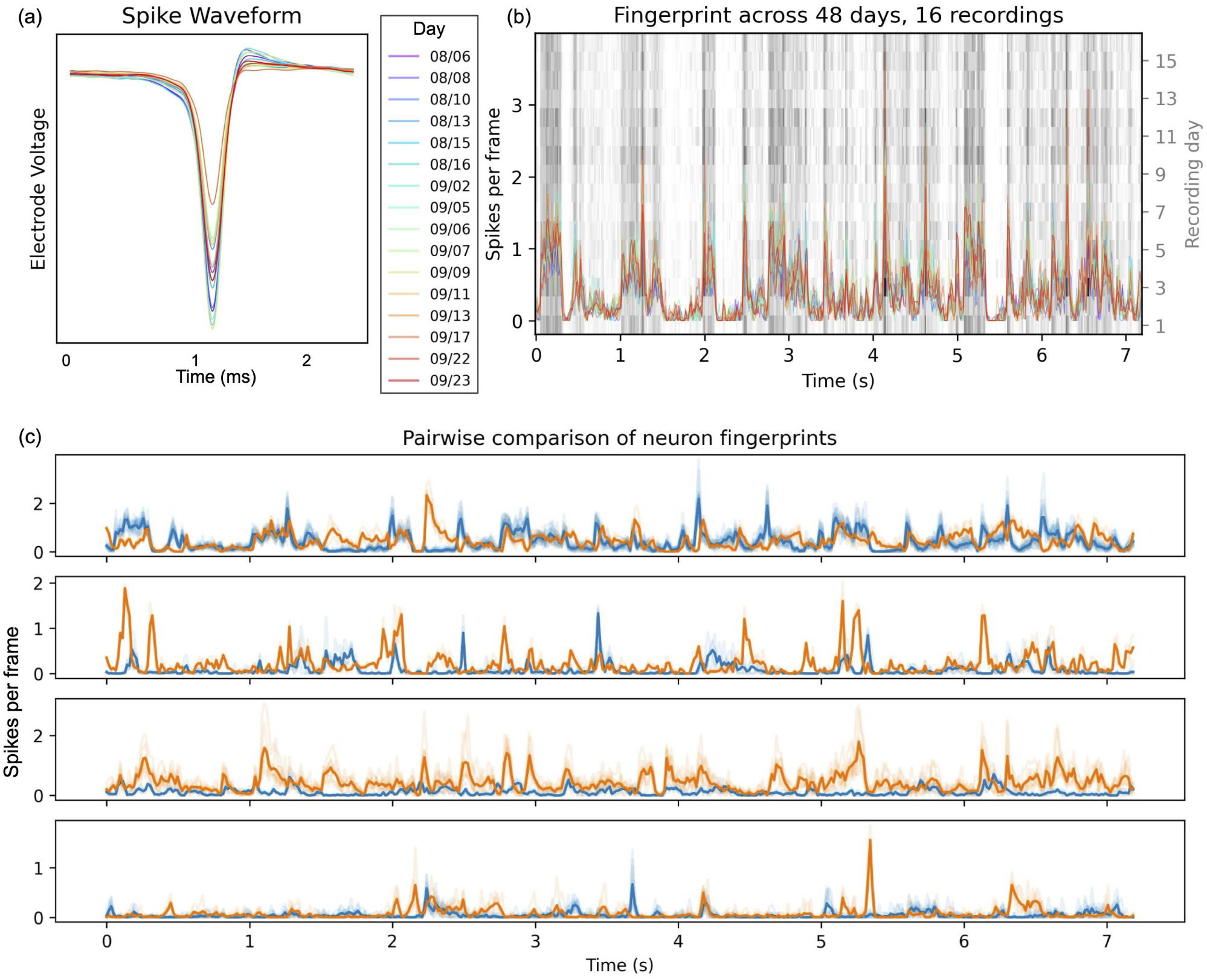
Tracking single neurons over multiple days. **(a)** The spike waveform of a V4 neuron, slightly varying across days. Because the spike waveform is unique to each neuron, a spike-sorting algorithm is used to group all spikes originating from a single neuron. However, the spike waveform may slightly change across days, due to small shifts of the electrode. Thus, spike-sorting algorithms are not suitable to track neurons over multiple days. **(b)** Peri-stimulus time histogram of a V4 neuron, recorded each day in response to a fixed stimulus. Because the response to the fixed stimulus is unique to each V4 neuron, the response is used as a functional fingerprint of the neuron. Recordings from different days are then identified as originating from the same neuron if the correlation between their functional fingerprints is high. This neuron’s fingerprint is consistent across 16 different recordings over a span of 48 days. The fingerprint for each day is plotted as a colored line, with the color legend shown on the right under “Day” with the corresponding recording dates. The 16 fingerprints are also shown as raster plots in the background, where each row is a recording day. This also shows how consistent a neuron’s fingerprint is over time and how strikingly time-locked they are to the stimulus. **(c)** Pairwise comparisons between fingerprints of two neurons. The fingerprints for eight neurons were compared, to assert that neuron fingerprints were not all driven by the fingerprint stimulus in a similar way. Each subplot compares a pair of 8 total neurons (one neuron in blue, the other in orange). The dark blue and orange traces are the means of the respective neurons’ fingerprints across days. Here it can be seen that the fingerprint of each neuron is quite different and uniquely driven by the stimulus.

In order to track neurons over multiple days, we used the peri-stimulus time histogram elicited by a naturalistic movie, as a functional fingerprint for each neuron. Each day, the recording session started with the same 10-second stimulus repeated 10 times, before proceeding to the main experiment. A spike-sorting algorithm was then used separately on each recording session, to group spikes originating from the same neuron. Spikes from different days were then matched, being identified as originating from the same neuron if the correlation between their fingerprints was high, typically 0.7 or greater. Fingerprints from each consecutive day were processed similarly, allowing us to track single neurons over extended periods of days or even weeks (see **Figure S1b**). In some instances, we were able to monitor a single V4 neuron for over 60 days.

Note that spike sorting methods are prone to both positive and negative error: a single cell might be treated as two different cells if the spike waveform changes over time, and multiple cells might be treated as a single cell if the two cells have similar spike waveforms. By augmenting information about similarity of spike waveform with information about similarity of tuning, we increase the sensitivity and accuracy of spike sorting. This occurs for two reasons. First, the dimensionality of the fingerprint response is much higher than the dimensionality of the spike waveform. Second, the variability in functional tuning of V4 neurons under naturalistic conditions is far larger than the variability in spike waveforms of neurons. This method is similar to cell identification methods already used and published by other labs ^62^. Thus, we are not the only group to have found this method to be useful.

Consistency of the receptive field estimate, using either synthetic stimuli or a pretrained DNN Because each V4 neuron has a limited spatial receptive field, information outside the receptive field is irrelevant for predicting the neuronal responses. Using this irrelevant information reduces the amount of relevant pixel information accessible to the model, thus leading to worse performance. Therefore, to maximize training efficiency, the HCE model was trained on only the portion of the stimulus that overlapped with the neuron’s receptive field. For each V4 neuron, the receptive field location was estimated, and the stimulus was cropped around the receptive field estimate. Estimating the spatial receptive field location of the V4 neurons was done using two different methods for the two animals.

**Figure S2:**
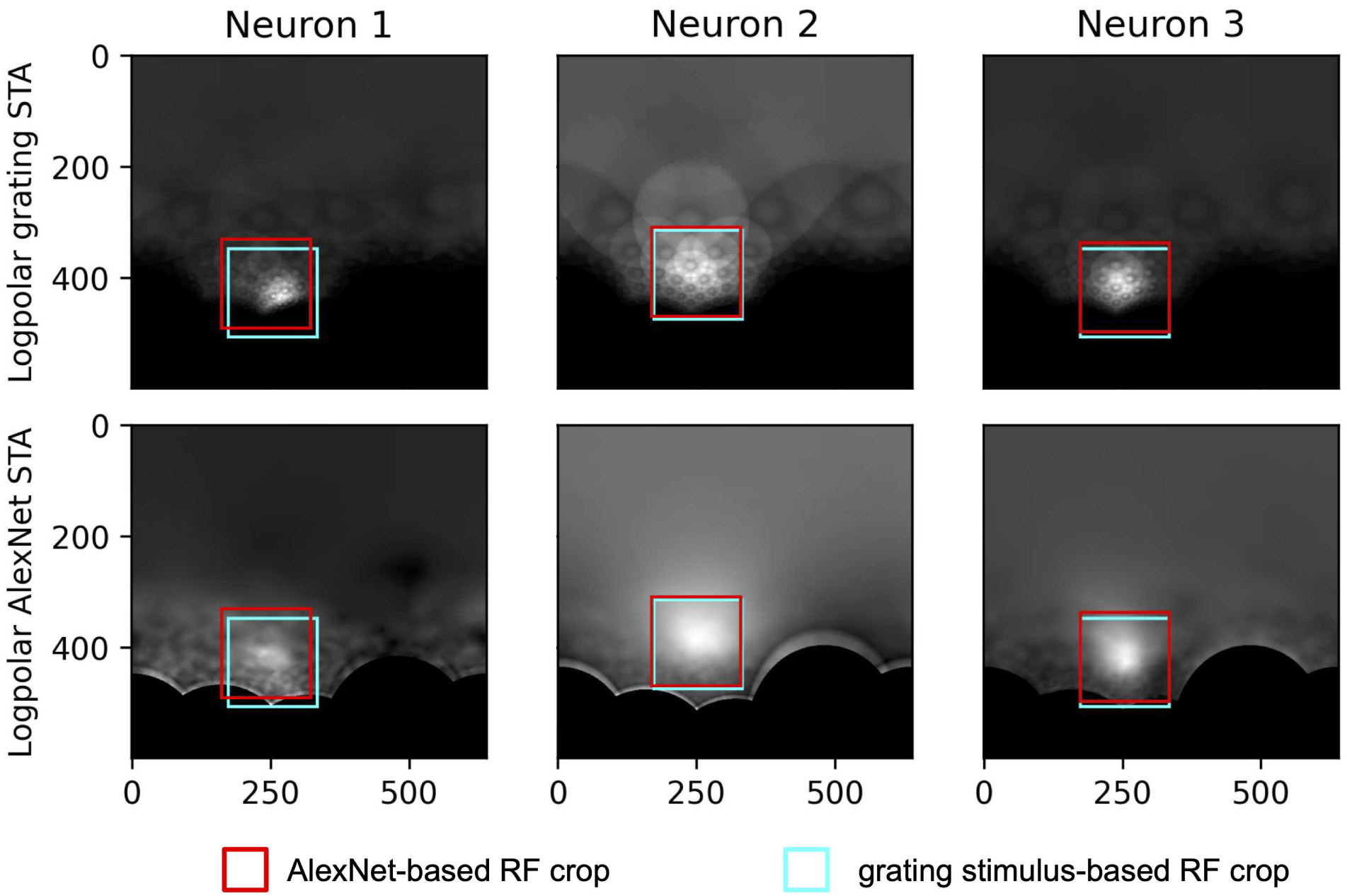
Comparing two methods to estimate V4 neurons’ receptive field locations. Two methods were used to estimate the receptive field locations for V4 neurons. The first method consisted of spike-triggered averaging (STA) over a set of synthetic grating stimuli. The second method consisted of sending the naturalistic stimuli through a pretrained AlexNet model, extracting the resulting feature maps, and combining them into spike-triggered averages. Both methods were applied on neurons from the first animal, and the two receptive field estimates were compared. The figure presents this comparison for three neurons, with the grating-based STA in the first row and the Alexnet-based STA in the second row. The receptive field estimates are shown using squares, with the Alexnet-based estimate in red, and the grating-based estimate in cyan. Both methods produced comparable estimates of the receptive field location.

For the first animal, an independent dataset was collected for estimating the receptive field location of each neuron. This independent dataset was composed of neuronal responses to synthetic stimuli such as Cartesian and non-Cartesian gratings ^10,11^ and parametric shapes ^12^. The synthetic stimuli were flashed for 83 ms at random locations while the animal fixated. The spike-triggered average of the responses to these stimuli was then computed by averaging all the stimulus frames that triggered a spike from the V4 neuron. The spike time series was temporally lagged to account for the neural response delay time relative to the movie frames, and the spatial receptive field was estimated on the temporal delay with the largest spike-triggered average response. In log-polar space, a region of 160×160 pixels was then manually selected around the area with the largest spike-triggered average. Projecting to log-polar space has the effect of normalizing the receptive field size of different neurons (see **Figure 1b**), and so the region was set to the same size for all neurons. The 160×160 pixel images were then downsampled to 64×64 pixels for computational efficiency, and used as input for the V1 module of the HCE model.

For the second animal, neuronal responses to the synthetic stimuli were not available for all V4 neurons. Therefore, a second method was used to estimate the spatial receptive fields. First, the stimulus movies were preprocessed using the pretrained AlexNet model ^68^, transforming each frame into abstract visual feature representations. Then, a spike-triggered average was computed across AlexNet’s feature activation maps to estimate the spatial receptive field of each neuron. A separate spike-triggered average was computed for each layer of the Alexnet architecture, and the best convolutional layer was selected as detailed below. Then, the activation map was log-polar-transformed with respect to fixation, and smoothed with a Gaussian filter with a standard deviation of 10 pixels. Finally, the center of mass of the smoothed activation map was computed and used to define the center of the receptive field, a region of 160×160 pixels. As with the first animal, the spike time series was temporally lagged to account for the neural response delay time relative to the movie frames, and the spatial receptive field was estimated on the temporal delay with the largest spike-triggered average response.

To test the consistency of the two methods, we applied both of them to neurons from the first animal who saw the synthetic stimuli (see **Figure S2**). We then compared the receptive field estimates. Out of all AlexNet layers, the fourth convolutional layer provided receptive field estimates most consistent with receptive fields estimated using the synthetic stimuli. Therefore, the fourth convolutional layer was used to estimate receptive fields for each V4 neuron in the second animal.

Comparison of the HCE model to an alternative model based on a pre-trained deep neural network

**Figure S3:**
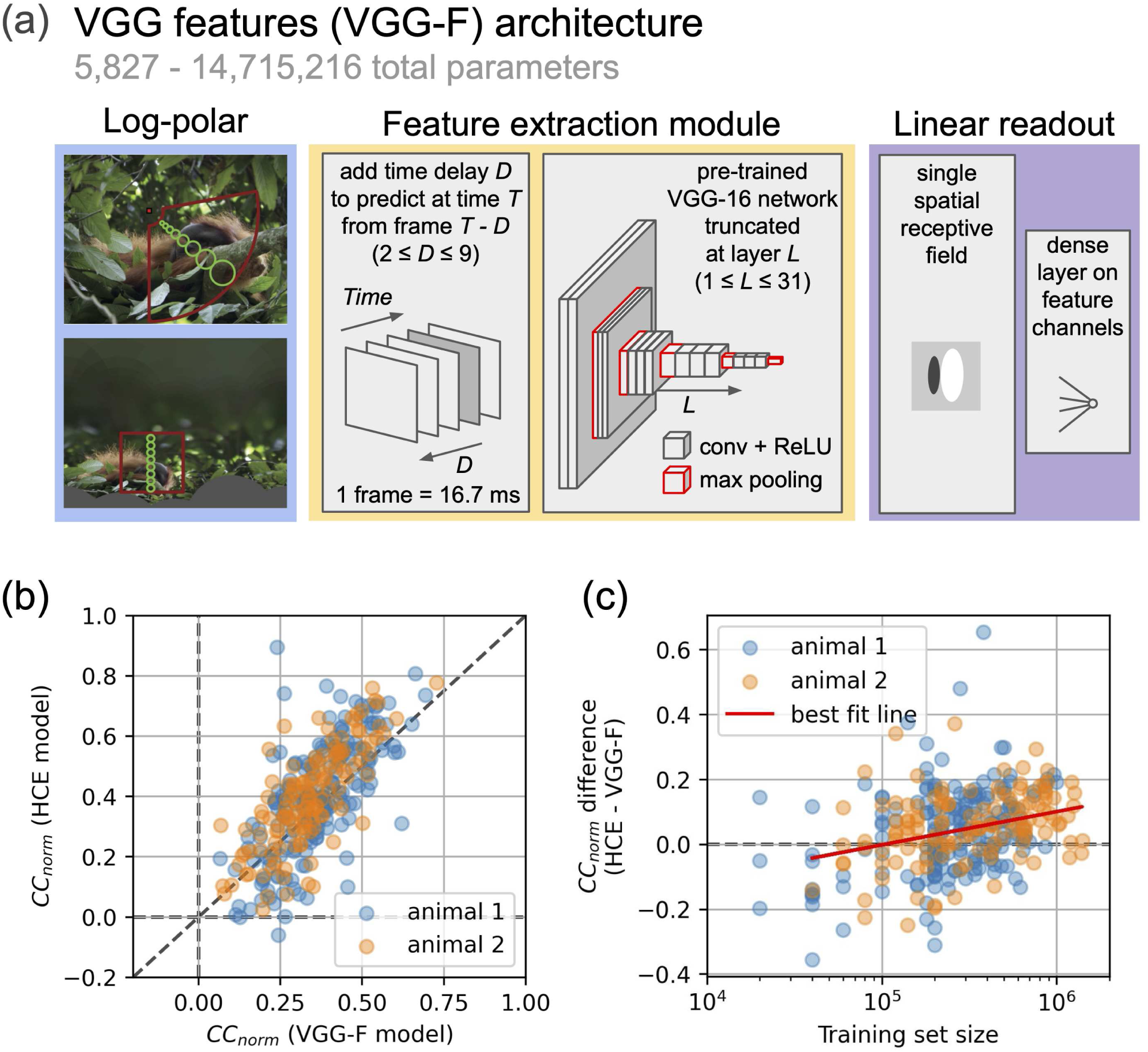
Comparison of the HCE architecture to a pre-trained deep neural network architecture (VGG-F). **(a)** Diagram of the VGG-F architecture. The VGG-F architecture leverages features from the VGG16 artificial neural network, a network trained to perform an object classification task. First, stimulus frames are delayed by *D* frames, and fed into a pretrained VGG16 network. Then, activations at layer *L* are extracted, and spatially combined with a learned linear receptive field. Finally, a learned linear layer combines the feature channels to produce a V4 neuron firing rate prediction. A separate VGG-F model was fit for every combination of layer *L* and delay *D*, and the best predicting model was selected based on predictive accuracy on a specific model-selection data set. **(b)** Scatter plot of the difference between HCE and VGG-F model performances as a function of the number of training frames that were presented to each V4 neuron. Neurons that are better predicted by the HCE model are above the horizontal dashed line. As the number of training stimuli increases, the difference in prediction accuracy of the HCE and the VGG-F model also increases, to the advantage of the HCE model (cross-validated r=0.10, regression fit). **(c)** Scatter plot of model performance (CC_norm_) for the HCE model (horizontal axis) and VGG-F model (vertical axis), on each individual V4 neuron of both animals. The HCE model consistently performs better than the VGG-F model on the V4 neuron population (p < 0.025, permutation test), particularly on well-predicted neurons.

Several previous studies have used a pre-trained deep neural network (DNN) to model responses of visual neurons ^61–64^. To compare the performance of the HCE model against this alternative, we used features provided by a standard DNN trained on an image classification task (VGG-16 ^84^; see **Methods**) to fit a separate model to the data recorded from each of the 313 V4 neurons in our sample. Prediction accuracy for the HCE model and the alternative VGG Features model (VGG-F) is shown in **Figure S3b**. The HCE model performs significantly better than the VGG-F model overall (p < 0.025, permutation test). The difference in performance between the two models is especially clear in the subset of neurons for which the largest set of training data are available (see **Figure S3c**). The HCE model produces better predictions than the VGG-F model for neurons with larger training data sets. This suggests that training the HCE model end-to-end, paired with large data sets, improves prediction performance compared to using transfer learning with pre-trained, task-based models. Thus, the HCE model outperforms a state-of-the-art DNN, while requiring far fewer parameters and while maximizing biological interpretability.

### Comparison of tuning properties across the two animals

The neurophysiology data reported in this paper were obtained from two different animals. The first animal yielded 183 neurons and the second yielded 130 neurons. To ensure that the HCE architecture presented here was not overfit, all development of the HCE architecture was performed using data from a subset of neurons from the first animal, and all of the data from the second animal were held out as a separate validation data set. To check that this procedure worked, we compared all receptive field tuning properties estimated throughout this paper across the two animals. The distributions of POPs were remarkably similar across the two animals when examining principal components (**Figures 5a-6a-7a**), color diagonal projections (**Figure 5c**), and effective ranks (**Figure 7b**).

The only notable difference that we were able to identify between the two animals occurred along the second principal component (PC) of the temporal profile embedding (**Figure 6a**). Inspection of the neurons projecting on both extremes of this PC indicates that it separates neurons that peak at time lags 2 or 3 (33 and 50 ms) from those that peak at time lags 4 or 5 (67 and 83 ms; **Figure 6a2**). To better quantify this difference, the temporal response profile of each neuron was approximated with continuous Gaussian windows parameterized by their center, width, and scale. Gaussian windows were fit to the temporal profile over an interval of five lags around the peak response lag. Considering only V4 neurons whose peak response lag is less than 100 ms, the histogram of Gaussian centers reveals that the median neuron peaks at 71 ms in the first animal and 62 ms in the second animal (p < 10^−12^, permutation test; **Figure 6c**). We suspect that this difference of 9 ms might be caused either by differences in the eccentricity of the recording grids, or by the difference in sizes or ages of the two animals. Thus, we conclude that V4 neurons sampled from these two animals are largely indistinguishable, excepting some minor difference in temporal conduction speed along the pathway from the eye to V4.

### How many predicted optimal patterns (POPs) are there for a single V4 neuron?

The population analysis described in the paper begins with the assumption that a POP accurately summarizes the tuning properties of a single neuron. This assumption will be valid only if a neuron has a single local optimum after stimulus optimization. However, if a neuron has multiple optima, then its tuning properties might not be summarized with a single POP. To address this issue, we used the stimulus optimization framework described above to determine whether V4 neurons had a single optimum or multiple optima. For each V4 neuron, the stimulus optimization was initialized using 100 different stimulus segments (14 frame, 232 ms) chosen at random while ensuring that the chosen segments had elicited a wide range of responses (i.e., high, medium and low). This produced a sample of 100 POPs for each neuron, chosen at random from the distribution of all possible POPs.

To determine whether this distribution was unimodal or multimodal for each neuron, the spectral, chromatic, and temporal embeddings were first calculated for each of the 100 POPs obtained for each neuron (see **Methods**). A gap statistic ^177,178^ was used to determine whether the distribution of POPs for each neuron was unimodal. The gap statistic is computed on a series of k-means clustering results. It compares the total sum of squared distances within each cluster to a uniform distribution. Specifically, if *W_k_* is the sum of the squared pairwise distances within each cluster *k,* the gap statistic *G_k_* is defined as the difference between the expected value of *W_k_,* denoted by *E* _[_*W_k_* _]_, and the actual value of *W_k_.* The optimal number of clusters for each neuron was determined by selecting the smallest value of *k* that satisfies the condition *G_k_ ≥G_k_*_+1_*−s_k_*_+1_, where *s_k_* _+1_ accounts for the standard deviation of the expected value estimate across multiple uniform distributions. This procedure selects the smallest *k* that maximizes the difference between the observed within-cluster distance and the expected within-cluster distance over a uniformly distributed sample. To compute the statistical significance of this result for each neuron in each embedding, the entire procedure was repeated 100 times by bootstrapping over each embedding. This produced 100 estimates of *G_k_* for each neuron and each possible clustering *k* . A permutation test was used to determine statistical significance, and this was corrected for the false discovery rate (FDR) ^161^.

Per the gap statistic, an optimal cluster number of *k = 1* resulted in the classification of a neuron as unimodal and *k > 1* as multimodal. In the spectral domain, 268 (85.6%) neurons were unimodal and 45 (14.4%) neurons were multimodal (p < 0.05 for 296 (94.6%) neurons, FDR corrected, permutation test). In the color domain, 263 (84.0%) neurons were unimodal and 50 (16.0%) neurons were multimodal (p < 0.05 for 273 (87.2%) neurons, FDR corrected, permutation test). In the time domain, 276 (88.2%) neurons were unimodal and 37 (11.8%) neurons were multimodal (p < 0.05 for 292 (93.3%) neurons, FDR corrected, permutation test). Thus, although the POPs synthesized for most V4 neurons did not depend on the video segment used for initialization, initial conditions did appear to affect POPs in 16% of the neurons.

Given that a substantial minority of V4 neurons have multimodal POPs, we next sought to determine the magnitude of the difference in tuning between the modes of multimodal POPs. If the difference in tuning between the modes of a multimodal POP is large relative to the difference in tuning between POPs estimated from different neurons, then the population-level analyses presented in this report might be biased. However, if all the modes of multimodal POPs have similar tuning in comparison to the distribution of POPs estimated on all neurons, then this effect can be ignored in subsequent population-level analysis. To quantify the variability of POPs within a neuron versus between neurons, the pairwise distances between POPs estimated within each multimodal V4 neuron were computed in the spatial-spectral embedding space, and these were compared to the pairwise distances between POPs between different V4 neurons. In all cases, the POPs estimated within single multimodal V4 neurons had a smaller pairwise distance than the POPs estimated between any two different neurons in our V4 sample (p < 0.05, permutation test; see **Figure S4e**). The same result was also obtained using the color-histogram embedding (see **Figure S4f**), and the time-profile embedding (see **Figure S4g**). Thus, although initial conditions affected the POPs estimated for 16% of the V4 neurons in our sample, this effect is small compared to the large differences in POPs obtained for different V4 neurons. Based on this finding, all subsequent population-level analyses focus on the single POP initialized with the most effective video sequence for each neuron.

**Figure S4:**
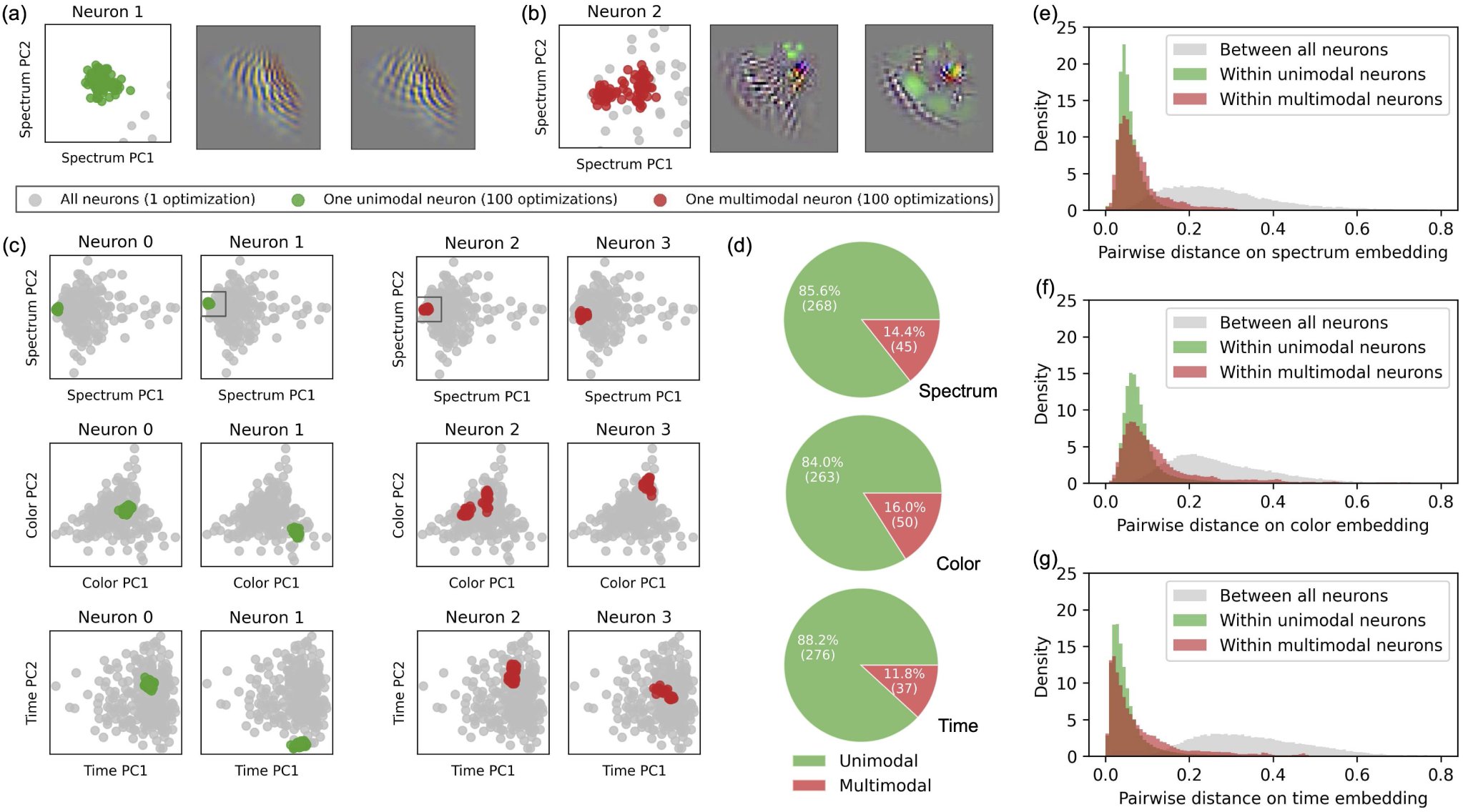
Stability of the POP generation procedure. To assess the stability of the POP generation procedure, a collection of 100 POPs was generated for each neuron, using 100 different stimulus segments for the procedure initialization. A clustering approach was then used to quantify the number of modes in the POP distribution of each neuron. **(a)** A unimodal neuron with a unimodal POP distribution. The distribution of POPs for a single “unimodal” neuron (shown in green) was projected onto the first two spatial-spectral PCs. The optimal POP recovered for each V4 neuron is shown in gray. The scatter plot is cropped to focus on the distribution of POPs for this single unimodal neuron. The distribution of POPs for this unimodal neuron is tightly clustered in this embedding space, indicating that they have similar tuning properties. Two example POPs from this unimodal neuron are shown to the right of the scatter plot. Even though the POPs were initialized with distinct stimuli, the two POPs are visually very similar. **(b)** A multimodal neuron with a multimodal POP distribution example. The distribution of POPs for a single “multimodal” neuron (shown in red) were projected onto the same embedding as in panel (a). The scatter plot is cropped to focus on the distribution of POP for this multimodal neuron as in part (a). When projected to the spectral embedding, the POPs for this multimodal neuron form two distinct clusters. Two example POPs from the multimodal neuron show that the different initializations result in visually distinct POPs that still share some common features. **(c)** Comparing the distribution of POPs for 4 single neurons with the distribution of POP of the entire V4 neuron population embedding. The distribution of POPs for a single neuron are shown projected onto the first two PCs of the spectral, chromatic, and temporal embeddings. The first two neurons have unimodal POP distributions (green), and the last two neurons have multimodal POP distribution (red). Unimodal neurons are plotted in green on the left, and multimodal neurons are plotted in red on the right. The boxes in the spectral plots for neuron 1 and neuron 2 correspond to the cropped scatter plots in (a) and (b) respectively. This shows that the variability across POPs within a single neuron is relatively small compared to the variability across POPs between all neurons. The boxes in the spectral plots for neuron 1 and neuron 2 correspond to the cropped scatter plots in (a) and (b) respectively. **(d)** Number of neurons classified as unimodal or multimodal based on the gap statistic computed on the spectrum, color, and time embeddings. In all three embeddings, less than one-sixth of the neurons were determined to be multimodal. **(e-g)** Density plots of pairwise distances on each of the three embeddings. The distribution of pairwise distances are plotted within the distributions of POPs for unimodal neurons (shown in green), within the distributions of POPs for multimodal neurons (shown in red), and across the optimal POPs for all V4 neurons (shown in gray). Within each embedding, the pairwise distance between the POPs of all V4 neurons is significantly greater than the pairwise distance within distributions of POPs in unimodal or multimodal neurons (p < 0.05, permutation test). Therefore, the variability of the POP generation procedure was considered small enough to not affect the rest of the population analysis described in our study.

### Relationship between space, color, and time PCs and model performance

For each PC in space, color, and time, the corresponding distribution of model performance was plotted. This was to determine whether there existed a trend in model performance corresponding to the PC. Neurons projected highly onto the first spatial PC tended to have lower model performance. Neurons that projected highly onto the first temporal PC also tended to have lower model performance. Interestingly, the second temporal PC, seems to separate neurons from the two animals based on model performance (see **Figure 6b-f**).

**Figure S5:**
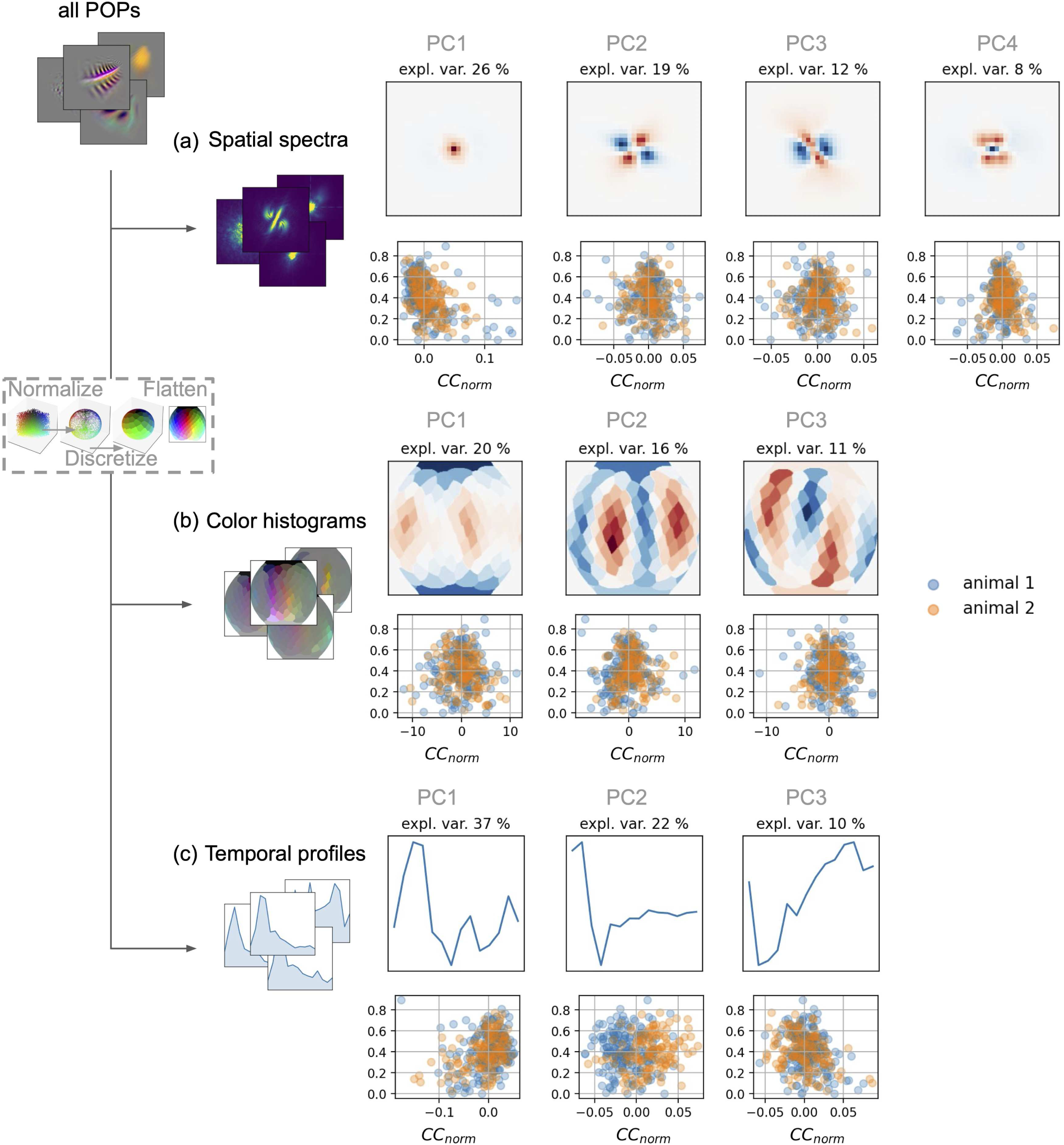
Relationship between space, color and time PCs and model performance. **(a)** The spatial spectra PCs from Figure 5 are visualized above scatter plots. The scatter plots correspond to each PC, where the y-axis shows the projection of each neuron onto that PC, and the x-axis is the model performance (denoted in CC_norm) of that neuron. The two animals are plotted in different colors to denote any animal-based differences between the V4 samples. Note that low-frequency neurons, demonstrated as neurons that project highly onto PC1, tend to have lower prediction accuracy. **(b)** The color histogram PCs from Figure 6 are visualized above scatter plots. **(c)** The temporal profile PCs from Figure 7 are visualized above scatter plots. Note that tonic neurons (neurons that project highly on PC1) tend to have lower prediction accuracy.

