## Supplementary materials for "A biologically-inspired hierarchical convolutional energy model predicts V4 responses to natural videos"

### Tracking neurons over days with spike sorting and fingerprinting

The construction of the large dataset employed in this study necessitated multi-day neuronal recording. Thus, spiking activity was recorded over many days across 96 electrodes of a Utah array. To identify individual neurons from the recordings, a spike-sorting algorithm was applied separately to each one of the 96 electrodes and for each day of recording (see **Methods**). The spike-sorting algorithm uses the fact that spikes from the same neuron have the same waveform, thereby grouping all spikes originating from a single neuron. However, the spike-sorting approach is not suitable to track neurons across days, because small shifts of the brain relative to the Utah array can have large effects on the spike waveforms (see **Figure S1a**). Consequently, an alternative strategy was adopted for multi-day neuron tracking.

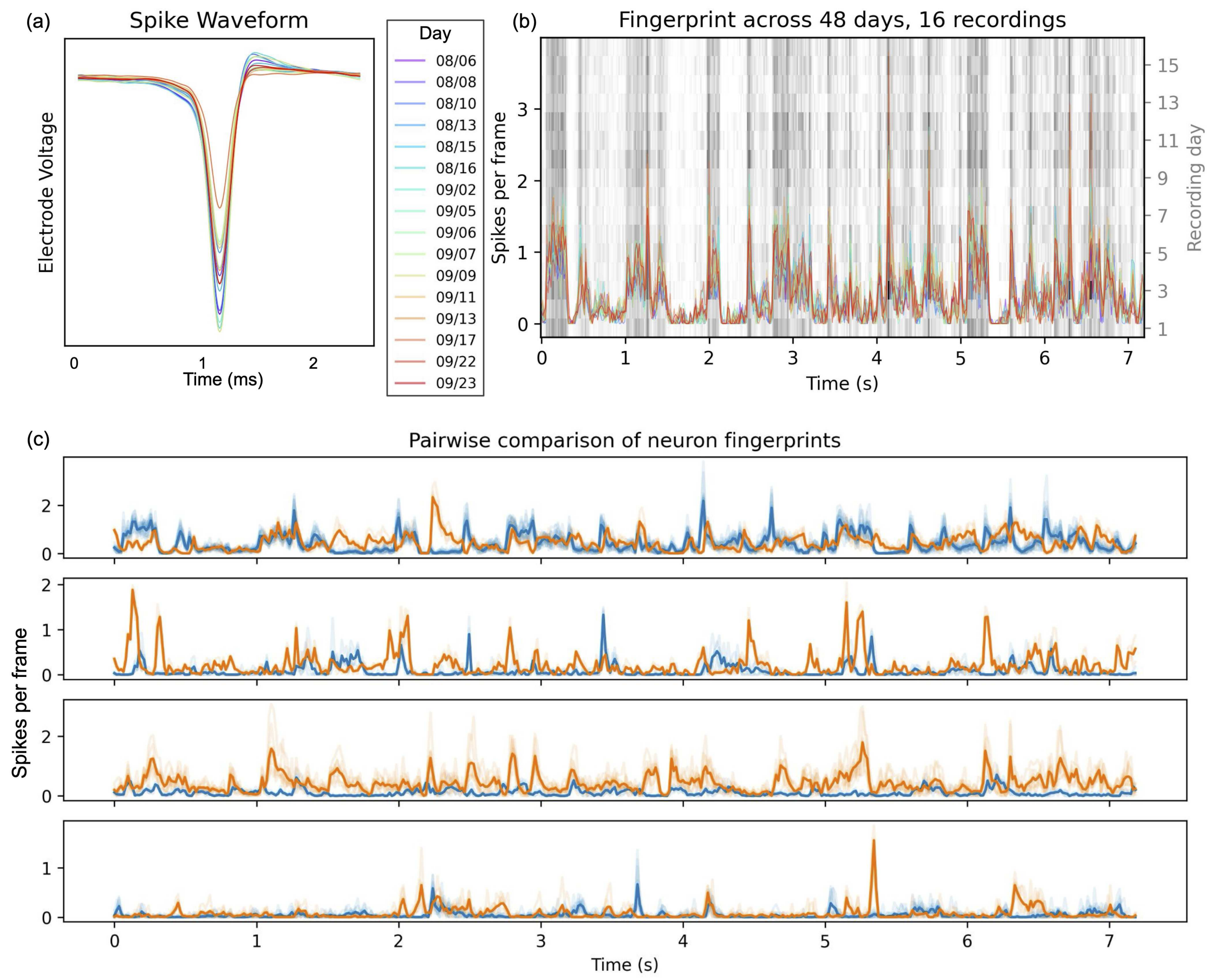


**Figure S1: Tracking single neurons over multiple days.** **(a)** The spike waveform of a V4 neuron, slightly varying across days. Because the spike waveform is unique to each neuron, a spike-sorting algorithm is used to group all spikes originating from a single neuron. However, the spike waveform may slightly change across days, due to small shifts of the electrode. Thus, spike-sorting algorithms are not suitable to track neurons over multiple days. **(b)** Peri-stimulus time histogram of a V4 neuron, recorded each day in response to a fixed stimulus. Because the response to the fixed stimulus is unique to each V4 neuron, the response is used as a functional fingerprint of the neuron.

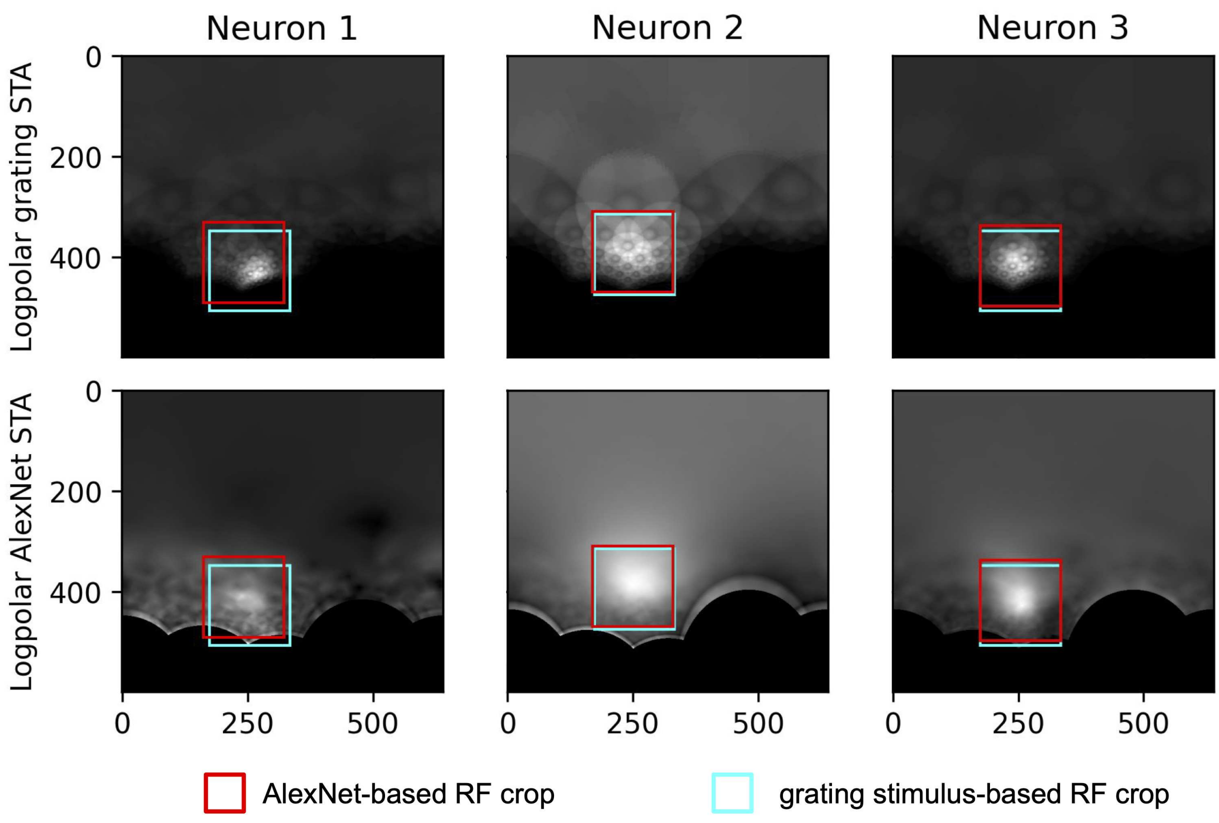


**Figure S2: Comparing two methods to estimate V4 neurons' receptive field locations.** Two methods were used to estimate the receptive field locations for V4 neurons. The first method consisted of spike-triggered averaging (STA) over a set of synthetic grating stimuli. The second method consisted of sending the naturalistic stimuli through a pretrained AlexNet model, extracting the resulting feature maps, and combining them into spike-triggered averages. Both methods were applied on neurons from the first animal, and the two receptive field estimates were compared. The figure presents this comparison for three neurons, with the grating-based STA in the first row and the Alexnet-based STA in the second row. The receptive field estimates are shown using squares, with the Alexnet-based estimate in red, and the grating-based estimate in cyan. Both methods produced comparable estimates of the receptive field location.

Comparison of the HCE model to an alternative model based on a pre-trained deep neural network

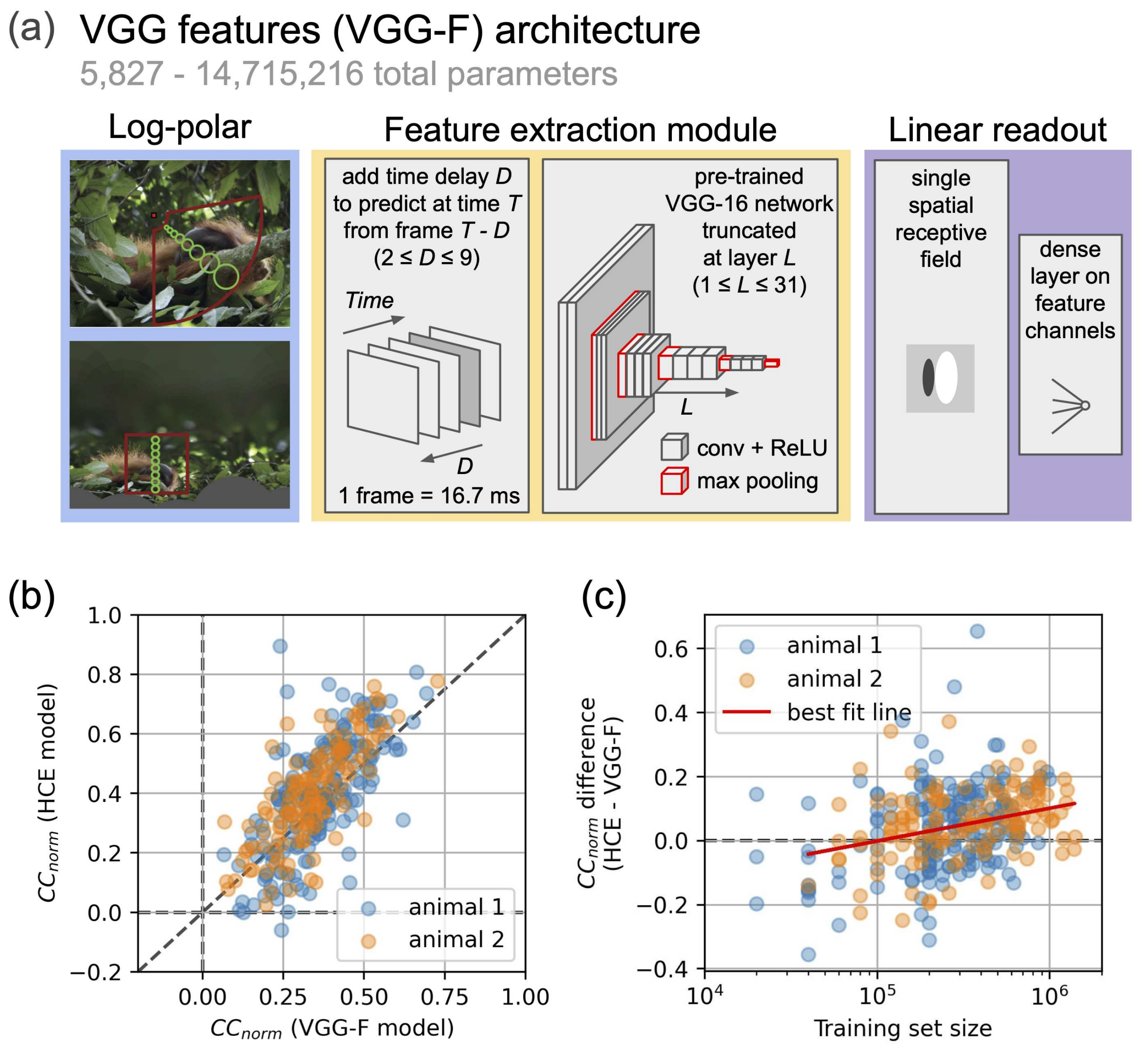


**Figure S3: Comparison of the HCE architecture to a pre-trained deep neural network architecture (VGG-F).** **(a)** Diagram of the VGG-F architecture. The VGG-F architecture leverages features from the VGG16 artificial neural network, a network trained to perform an object classification task. First, stimulus frames are delayed by *D* frames, and fed into a pretrained VGG16 network. Then, activations at layer *L* are extracted, and spatially combined with a learned linear receptive field. Finally, a learned linear layer combines the feature channels to produce a V4 neuron firing rate prediction. A separate VGG-F model was fit for every combination of layer *L* and delay *D*, and the best predicting model was selected based on predictive accuracy on a specific model-selection data set. **(b)** Scatter plot of the difference between HCE and VGG-F model performances as a function of the number of training frames that were presented to each V4 neuron. Neurons that are better predicted by the HCE model are above the horizontal dashed line. As the number of training stimuli increases, the difference in prediction accuracy of the HCE and the VGG-F model also increases, to the advantage of the HCE model (cross-validated r=0.10, regression fit). **(c)** Scatter plot of model performance ($CC_{norm}$) for the HCE model (horizontal axis) and VGG-F model (vertical axis), on each individual V4 neuron of both animals. The HCE model consistently performs better than the VGG-F model on the V4 neuron population ($p < 0.025$, permutation test), particularly on well-predicted neurons.

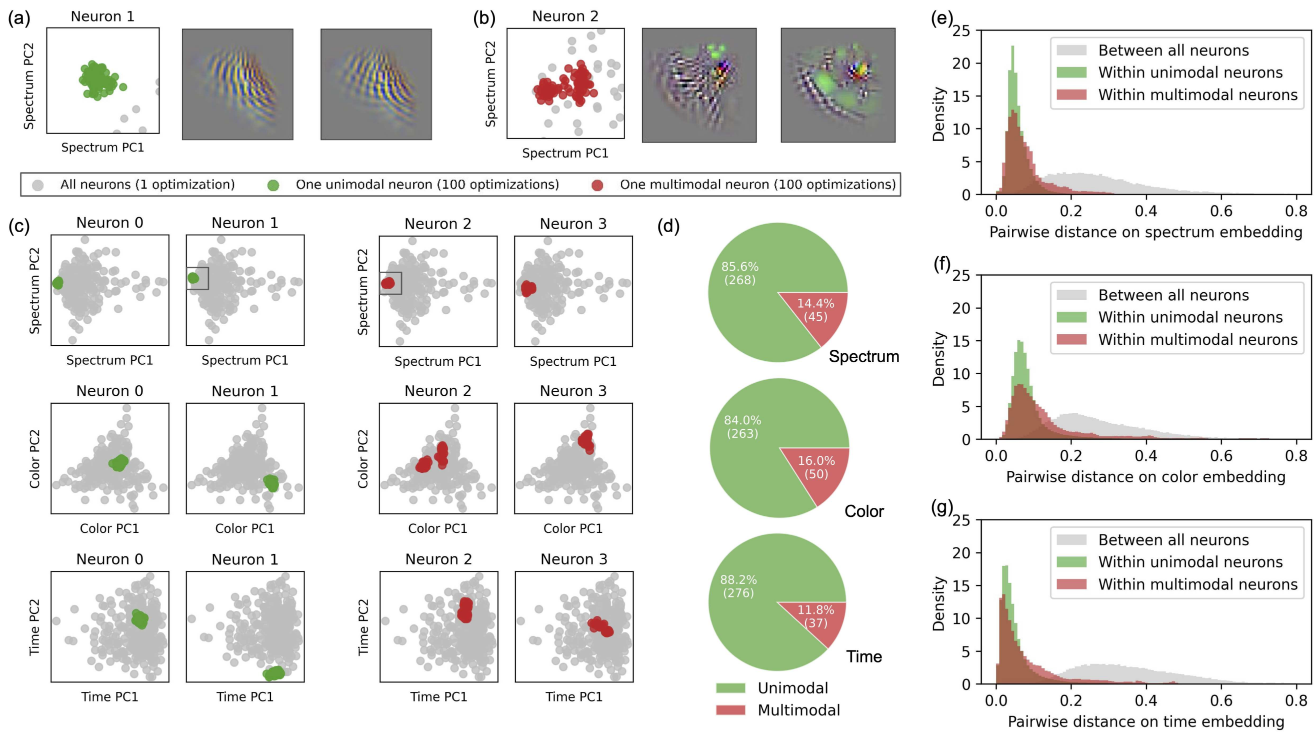


**<u>Figure S4: Stability of the POP generation procedure.</u>** To assess the stability of the POP generation procedure, a collection of 100 POPs was generated for each neuron, using 100 different stimulus segments for the procedure initialization. A clustering approach was then used to quantify the number of modes in the POP distribution of each neuron. **(a)** A unimodal neuron with a unimodal POP distribution. The distribution of POPs for a single "unimodal" neuron (shown in green) was projected onto the first two spatial-spectral PCs. The optimal POP recovered for each V4 neuron is shown in gray. The scatter plot is cropped to focus on the distribution of POPs for this single unimodal neuron. The distribution of POPs for this unimodal neuron is tightly clustered in this embedding space, indicating that they have similar tuning properties. Two example POPs from this unimodal neuron are shown to the right of the scatter plot. Even though the POPs were initialized with distinct stimuli, the two POPs are visually very similar. **(b)** A multimodal neuron with a multimodal POP distribution example. The distribution of POPs for a single "multimodal" neuron (shown in red) were projected onto the same embedding as in panel (a). The scatter plot is cropped to focus on the distribution of POP for this multimodal neuron as in part (a). When projected to the spectral embedding, the POPs for this multimodal neuron form two distinct clusters. Two example POPs from the multimodal neuron show that the different initializations result in visually distinct POPs that still share some common features. **(c)** Comparing the distribution of POPs for 4 single neurons with the distribution of POP of the entire V4 neuron population embedding. The distribution of POPs for a single neuron are shown projected onto the first two PCs of the spectral, chromatic, and temporal embeddings. The first two neurons have unimodal POP distributions (green), and the last two neurons have multimodal POP distribution (red). Unimodal neurons are plotted in green on the left, and multimodal neurons are plotted in red on the right. The boxes in the spectral plots for neuron 1 and neuron 2 correspond to the cropped scatter plots in (a) and (b) respectively. This shows that the variability across POPs within a single neuron is relatively small compared to the variability across POPs between all neurons. The boxes in the spectral plots for neuron 1 and neuron 2 correspond to the cropped scatter plots in (a) and (b) respectively. **(d)** Number of neurons classified as unimodal or multimodal based on the gap statistic computed on the spectrum, color, and time embeddings. In all three embeddings, less than one-sixth of the neurons were determined to be multimodal. **(e-g)** Density plots of pairwise distances on each of the three embeddings. The distribution of pairwise distances are plotted within the distributions of POPs for unimodal neurons (shown in green), within the distributions of POPs for multimodal neurons (shown in red), and across the optimal POPs for all V4 neurons (shown in gray). Within each embedding, the pairwise distance between the POPs of all V4 neurons is significantly greater than the pairwise distance within distributions of POPs in unimodal or multimodal neurons ($p < 0.05$, permutation test). Therefore, the variability of the POP generation procedure was considered small enough to not affect the rest of the population analysis described in our study.

**Relationship between space, color, and time PCs and model performance**

For each PC in space, color, and time, the corresponding distribution of model performance was plotted. This was to determine whether there existed a trend in model performance corresponding to the PC. Neurons projected highly onto the first spatial PC tended to have lower model performance. Neurons that projected highly onto the first temporal PC also tended to have lower model performance. Interestingly, the second temporal PC, seems to separate neurons from the two animals based on model performance (see **Figure 6b-f**).

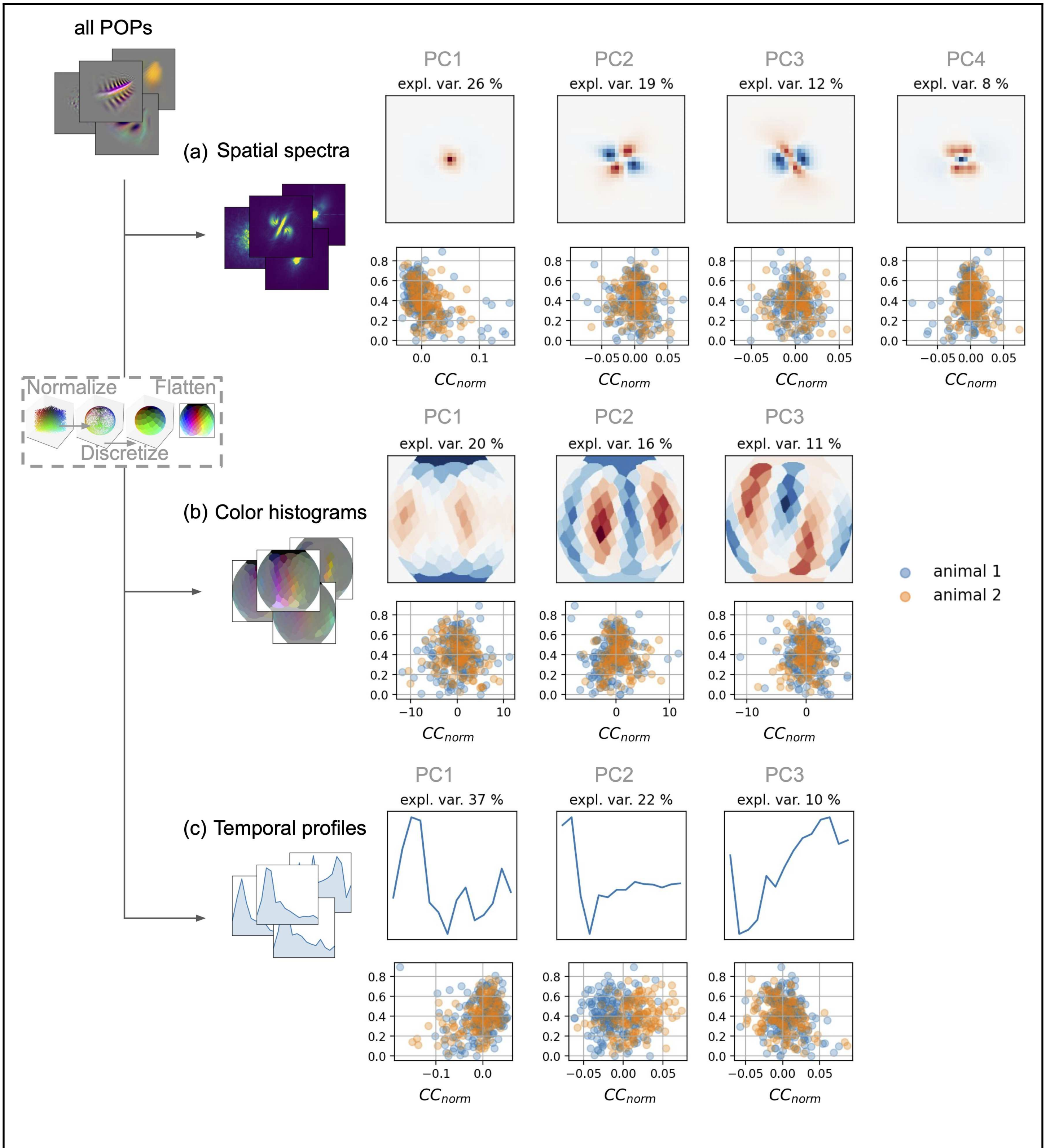


**Figure S5: Relationship between space, color and time PCs and model performance.**
**(a)** The spatial spectra PCs from Figure 5 are visualized above scatter plots. The scatter plots correspond to each PC, where the y-axis shows the projection of each neuron onto that PC, and the x-axis is the model performance (denoted in CC_norm) of that neuron. The two animals are plotted in different colors to denote any animal-based differences between the V4 samples. Note that low-frequency neurons, demonstrated as neurons that project highly onto PC1, tend to have lower prediction accuracy. **(b)** The color histogram PCs

from Figure 6 are visualized above scatter plots. **(c)** The temporal profile PCs from Figure 7 are visualized above scatter plots. Note that tonic neurons (neurons that project highly on PC1) tend to have lower prediction accuracy.

144. Srivastava, N., Hinton, G., Krizhevsky, A., Sutskever, I. & Salakhutdinov, R. Dropout: A simple way to prevent neural networks from overfitting. https://www.jmlr.org/papers/volume15/srivastava14a/srivastava14a.pdf?utm_content=buffer79b43&utm_medium=social&utm_source=twitter.com&utm_campaign=buffer,.

145. Schmolesky, M. T. *et al.* Signal timing across the macaque visual system. *J. Neurophysiol.* **79**, 3272–3278 (1998).

146. Zheng, S. & Kwok, J. T. Follow the Moving Leader in Deep Learning. **70**, 4110–4119 (2017).

147. Kingma, D. P. & Ba, J. Adam: A Method for Stochastic Optimization. *arXiv [cs.LG]* (2014).

157. Efron, B. & Tibshirani, R. J. *An Introduction to the Bootstrap*. (CRC Press, 1994).

158. Lebart, L., Morineau, A. & Piron, M. *Statistique Exploratoire Multidimensionnelle*. vol. 3 (Dunod Paris, 1995).

159. González, Á. Measurement of Areas on a Sphere Using Fibonacci and Latitude–Longitude Lattices. *Math. Geosci.* **42**, 49 (2009).

160. Robinson, A. H. A new map projection: Its development and characteristics. *International yearbook of cartography* **14**, 145–155 (1974).

161. Benjamini, Y. & Hochberg, Y. Controlling the false discovery rate: A practical and powerful approach to multiple testing. *J. R. Stat. Soc.* **57**, 289–300 (1995).

162. Chollet, F. & Others. Keras: Deep learning library for theano and tensorflow. *URL: https://keras. io/k*.

163. The Theano Development Team *et al.* Theano: A Python framework for fast computation of mathematical expressions. *arXiv [cs.SC]* (2016).

164. Paszke, A. *et al.* PyTorch: An imperative style, high-performance deep learning library. in *Advances in Neural Information Processing Systems* 8024–8035 (2019).

165. Bradski, G. The openCV library. *Dr. Dobb's Journal: Software Tools for the Professional Programmer* **25**, 120–123 (2000).

166. Klein, A. *et al. Imageio/imageio: v2.28.1*. (2023). doi:10.5281/zenodo.7882642.

167. Clark, J. A. (alex). *Pillow (PIL Fork) Documentation*. (2023).

168. Collette, A. *Python and Hdf5: Unlocking Scientific Data*. (O'Reilly Media, 2013).

178. Mohajer, M., Englmeier, K.-H. & Schmid, V. J. A comparison of Gap statistic definitions with and without logarithm function. *arXiv [stat.ME]* (2011).
